# HSPB1 facilitates chemoresistance through inhibiting ferroptotic cancer cell death and regulating NF-κB signaling pathway in breast cancer

**DOI:** 10.1101/2022.10.25.513668

**Authors:** Yiran Liang, Yajie Wang, Yan Zhang, Fangzhou Ye, Dan Luo, Yaming Li, Yuhan Jin, Dianwen Han, Zekun Wang, Bing Chen, Wenjing Zhao, Lijuan Wang, Xi Chen, Tingting Ma, Xiaoli Kong, Qifeng Yang

**Author notes:** These authors contributed equally to this work (Yiran Liang, Yajie Wang). Correspondence: Qifeng Yang, Qilu Hospital, Cheeloo College of Medicine, Shandong University, Wenhua Xi Road No. 107, Jinan 250012, Shandong, China.

## Abstract

Chemoresistance is one of the major causes of therapeutic failure and poor prognosis for breast cancer patients, especially for triple-negative breast cancer patients. However, the underlying mechanism remains elusive. Here, we identified novel functional roles of heat shock protein beta-1 (HSPB1), regulating the chemoresistance and ferroptotic cell death in breast cancer. Based on TCGA and GEO databases, HSPB1 expression was upregulated in breast cancer tissues and associated with poor prognosis of breast cancer patients, which was considered as an independent prognostic factor for breast cancer. Functional assays revealed that HSPB1 could promote cancer growth and metastasis in vitro and in vivo. Furthermore, HSPB1 facilitated doxorubicin resistance through protecting breast cancer cells from drug-induced ferroptosis. Mechanistically, HSPB1 could bind with Ikβ-α and promote its ubiquitination-mediated degradation, leading to increased nuclear translocation and activation of NF-κB signaling. In addition, HSPB1 overexpression led to enhanced secretion of IL6, which further facilitated breast cancer progression. These findings revealed that HSPB1 upregulation might be a key driver to progression and chemoresistance through regulating ferroptosis in breast cancer, while targeting HSPB1 could be an effective strategy against breast cancer.

## Introduction

Breast cancer is one of the most prevalent malignancies in women, accounting for more than 24% of new female cancer cases and about 15% of cancer-related death around the world (1). Based on the expression status of estrogen receptor (ER), progesterone receptor (PR), and human epidermal growth factor receptor 2 (HER2), breast cancer is generally classified in to four subtypes (2): Luminal A, Luminal B, HER2-enriched, or triple-negative breast cancer (TNBC). TNBC is a subtype of breast cancer characterized by the absence of ER and PR and the lack of HER2 amplification or overexpression (3), accounting for 15-20% of all invasive breast cancers (4). Due to the absence of druggable molecular drivers in TNBC, chemotherapy is still the mainstay of systemic treatment for TNBC patents (5). Nevertheless, intrinsic and acquired drug resistance greatly limited the efficiency of chemotherapy, leading to high rates of metastasis and poor prognosis in breast cancer patients (6). Therefore, characterization of the underlying molecular mechanisms of chemoresistance would help to develop novel therapeutic strategy to enhance the efficacy of chemotherapy in breast cancer patients.

Numerous mechanisms have been reported to be responsible for chemoresistance (7–9), including enhanced efflux of intracellular drugs, epithelial-to-mesenchymal transition (EMT), improved DNA damage repair, and increased cell death inhibition. Ferroptosis, a recently recognized form of cell death, is mainly caused by excessive iron-dependent lipid peroxidation and results in iron-mediated oxidative damage of cell membranes with an increasing level of intracellular reactive oxygen species (ROS) (10). It has unique characteristics distinct from apoptosis, necrosis, and autophagy in morphology, biochemistry and genetics (11, 12). Accumulated evidences indicated that ferroptosis played a significant role in the fate of cancer cells and response to various cancer treatments, such as chemotherapy, radiotherapy, and immunotherapy (13). Tumor cells with drug resistance and high metastatic tendency showed higher susceptibility to ferroptosis (14, 15), indicating that targeting the negative regulators of ferroptosis might further render chemoresistant cancer cells susceptible to ferroptotic cell death (16). In addition, damage-related molecular patterns (DAMPs) released from the ferroptotic cancer cells could enhance inflammation and immune responses through acting on surrounding cells (17–19). Various reagents, factors and drugs, such as RSL3, erastin, and sorafenib, have been demonstrated to inhibit tumor growth through inducing ferroptosis (20, 21). Therefore, further studies are required to elucidate the molecular mechanism of ferroptosis, which would facilitate the implementation of novel approaches to overcome chemoresistance.

In the current study, based on the differential analysis of data from TCGA and GEO databases, we identified that the expression of Heat shock protein beta-1 (HSPB1), a member of the small heat shock proteins, was significantly upregulated in breast cancer tissues. Moreover, high expression of HSPB1 was associated with poor prognosis of breast cancer patients. Functional experiments verified that HSPB1 showed a crucial role in the progression of breast cancer. We further revealed the relevance of HSPB1 in the regulation of chemoresistance of breast cancer for the first time, which was mediated by chemotherapeutics-induced ferroptosis. Our study highlights HSPB1 as a novel regulator of chemoresistance in breast cancer, providing the potential that targeting HSPB1 might be a novel strategy to prevent breast cancer progression and overcome therapy resistance.

## Results

### HSPB1 is elevated in breast cancer tissues, and high HSPB1 expression is associated with poor prognosis of breast cancer patients

We first analyzed the mRNA expression in several public breast cancer datasets, and hierarchical clustering analysis revealed that the expression of HSPB1 was upregulated in breast cancer tissues compared to normal tissues based on TCGA and GEO databases (Figure 1A, Figure 1-figure supplement 1A). Moreover, higher RNA expression of HSPB1 was discovered in several other cancers (Figure 1-figure supplement 1B), such as cervical and endocervical cancer (CESC), cholangiocarcinoma (CHOL), esophageal carcinoma (ESCA), glioblastoma (GBM), kidney renal papillary cell carcinoma (KIRP), and liver hepatocellular carcinoma (LIHC). We further detected the expression of HSPB1 in human breast cancer tissues and adjacent normal tissues in a cohort of patients from Qilu hospital, and upregulated mRNA expression of HSPB1 was also detected in breast cancer tissues (Figure 1B). Consistently, the IHC staining also revealed that the protein expression of HSPB1 was also upregulated in breast cancer tissues compared to normal tissues according to Qilu cohort and TCGA dataset (Figure 1C, Figure 1-figure supplement 1C). In addition, HSPB1 expression was more abundant in breast cancer cells than in the normal cells (Figure 1D-E), further supporting the tumor-promoting role of HSPB1 in breast cancer. We also investigate the clinical significance of HSPB1, and noted that elevated HSPB1 expression was associated with worse overall survival by analyzing several publicly available datasets (Figure 1-figure supplement 1D). Furthermore, Kaplan–Meier analysis showed that breast cancer patients with high HSPB1 expression had poorer overall survival and disease-free survival based on analysis of 131 cases of breast cancers (Figure 1F-G). Collectively, these findings indicated that HSPB1 expression was increased in breast cancer tissues and associated with poor prognosis of breast cancer patients.

**Figure 1.**
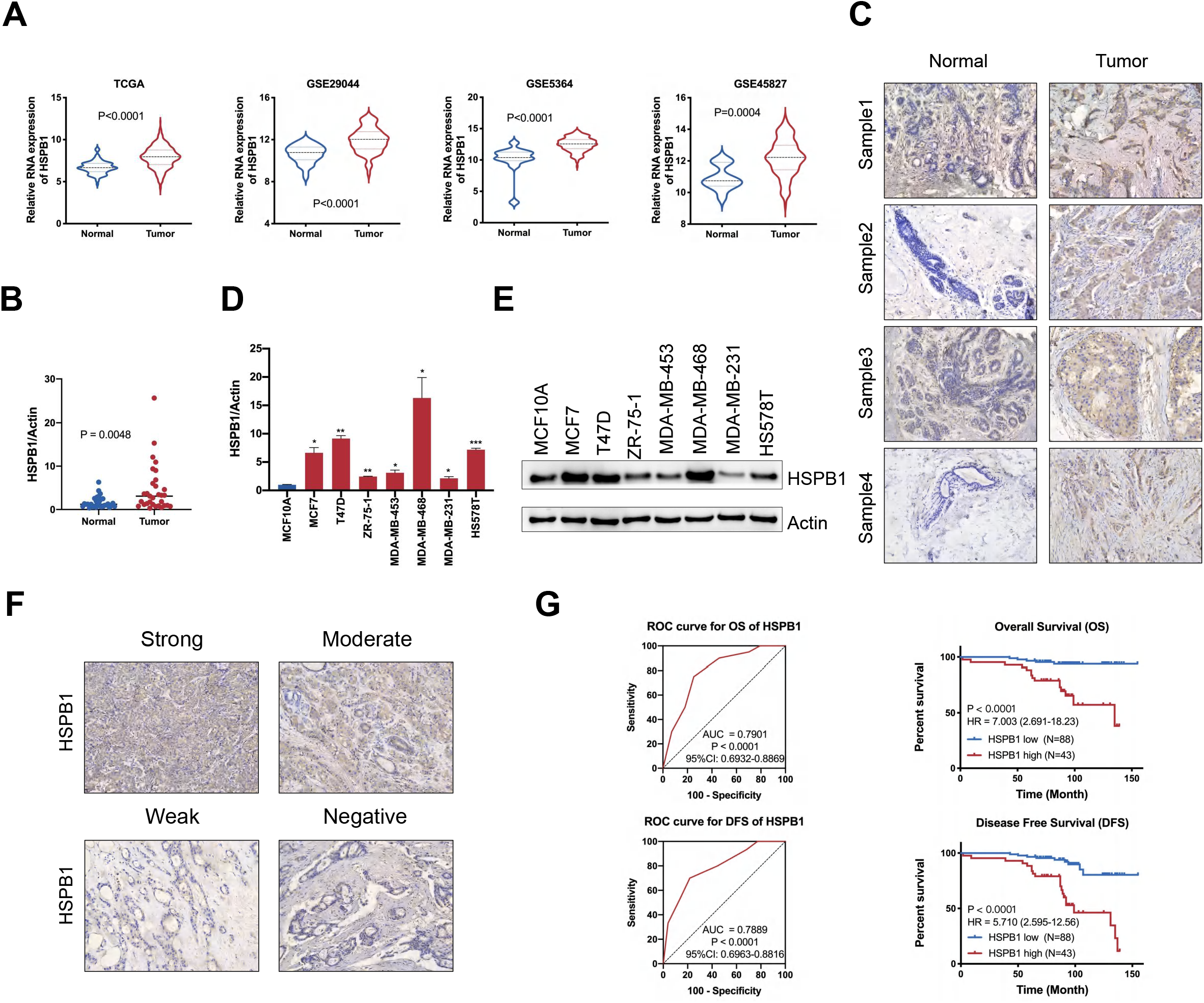
HSPB1 was associated with poor response to chemotherapy of breast cancer patients. (A) The expression of HSPB1 was elevated in breast cancer tissues compared to normal tissues according to TCGA and GEO database. (B-C) The RNA (B) and protein (C) expression of HSPB1 was upregulated in breast cancer tissues compared to normal tissues based on Qilu cohort. (D-E) The RNA (D) and protein (E) expression of HSPB1 was upregulated in breast cancer cells compared to normal cell. (F) HSPB1 expression was detected by IHC. (G) Patients with breast cancer were divided into two subsets with low and high HSPB1 expression based on ROC analysis. The correlation between HSPB1 expression and overall survival or disease-free survival in breast cancer was calculated with the log-rank test. (* P < 0.05, ** P < 0.01, *** P < 0.001) **Figure 1-source data 1** Original western blots for Figure 1E.

### Clinical significance of HSPB1 in breast cancer

The chi-square test was used to evaluated the correlations between HSPB1 expression and clinicopathologic features of breast cancer patients, such as age, tumor stage, LN metastasis, distant metastasis, histologic grade, ER status, PR status, HER-2 status, and Ki67 expression. The results indicated that high expression of HSPB1 was significantly associated with distant metastasis in breast cancer (p<0.001), indicating a potential role of HSPB1 in breast cancer progression (Table 1). Furthermore, univariate and multivariate Cox regression analyses were performed to screen the potential prognostic factors in breast cancer. We found that age, >3 LN metastasis, and HSPB1 expression were independent prognostic indicators for overall survival of breast cancer (Table 2). Significantly, age, LN metastasis, distant metastasis, and HSPB1 expression were considered as independent prognostic indicators for disease-free survival of breast cancer (Table 3). These results indicated that HSPB1 was an unfavorable prognostic biomarker in breast cancer.

**Table 1.** Correlations between HSPB1 expression and clinicopathologic features in 131 breast cancer patients.

| Characteristics | HSPB1-low(n=88) | HSPB1-high(n=43) | P value |
| --- | --- | --- | --- |
| Age |  |  | p=0.291 |
| <59 | 67(76.14%) | 29(67.44%) |  |
| ≥59 | 21(23.86%) | 14(32.56%) |  |
| Tumor stage |  |  | p=0.806 |
| ≤T1 | 44(50.00%) | 20(46.51%) |  |
| >T1 | 43(48.86%) | 23(53.49%) |  |
| unknown | 1(1.14%) | 0(0.00%) |  |
| LN metastasis |  |  | p=0.166 |
| 0 | 61(69.32%) | 23(53.48%) |  |
| 1-3 | 16(18.18%) | 10(23.26%) |  |
| >3 | 11(12.50%) | 10(23.26%) |  |
| Distant metastasis |  |  | **p<0.001** |
| No | 76(86.36%) | 21(48.84%) |  |
| Yes | 7(7.95%) | 6(13.95%) |  |
| unknown | 5(5.68%) | 16(37.21%) |  |
| Histologic grade |  |  | p=0.166 |
| G1 | 7(7.95%) | 0(0.00%) |  |
| G2 | 46(52.27%) | 20(46.51%) |  |
| G3 | 27(30.68%) | 19(44.19%) |  |
| unknown | 8(9.09%) | 4(9.30%) |  |
| ER status |  |  | p=0.721 |
| negative | 52(59.09%) | 24(55.81%) |  |
| positive | 36(40.91%) | 19(44.19%) |  |
| PR status |  |  | p=0.463 |
| negative | 55(62.50%) | 24(55.81%) |  |
| positive | 33(37.50%) | 19(44.19%) |  |
| HER-2 status |  |  | p=0.154 |
| negative | 81(92.05%) | 35(81.40%) |  |
| positive | 1(1.14%) | 2(4.65%) |  |
| unknown | 6(6.81%) | 6(13.95%) |  |
| Ki67 expression |  |  | p=0.052 |
| low | 22(25.00%) | 6(13.95%) |  |
| high | 66(75.00%) | 35(81.40%) |  |
| unknown | 0(0.00%) | 2(4.65%) |  |

Abbreviation: LN=lymph nodes; ER=estrogen receptor; PR=progesterone receptor; HER-2: human epidermal growth factor receptor-2; p value < 0.05 marked in bold font to show statistical significant.

**Table 2.** Univariate and multivariate Cox regression analyses for OS of 131 breast cancer patients.

|  | Univariate analysis |  | Multivariate analysis |  |
| --- | --- | --- | --- | --- |
|  | HR(95%CI) | p value | HR(95%CI) | p value |
| Age($\geq 59$ vs. $< 59$) | 3.028(1.259-7.281) | **0.013** | 2.704(1.123-6.510) | **0.026** |
| Histologic grade |  |  |  |  |
| G2 vs. G1 | 9317.972(0.000-3.001E+78) | 0.917 | - | - |
| G3 vs. G1 | 8650.070(0.000-2.787E+78) | 0.918 | - | - |
| Unknown vs. G1 | 8523.247(0.000-2.758E+78) | 0.918 | - | - |
| Tumor size($> T1$ vs. $\leq T1$+unknown) | 1.313(0.543-3.172) | 0.545 | - | - |
| LN metastasis |  |  |  |  |
| 1-3 vs. 0 | 1.983(0.580-6.780) | 0.275 | 2.067(0.590-7.241) | 0.256 |
| $> 3$ vs. 0 | 8.050(2.902-22.326) | **<0.001** | 6.213(2.165-17.827) | **0.001** |
| Distant metastasis (Yes vs. No+unknown) | 1.579(0.462-5.394) | 0.466 | - | - |
| ER status(pos vs. neg) | 1.017(0.400-2.584) | 0.973 | - | - |
| PR status(pos vs. neg) | 0.878(0.339-2.272) | 0.788 | - | - |
| HER-2 status(pos vs. neg+unknown) | 2.433(0.322-18.388) | 0.389 | - | - |
| Ki67 expression(pos vs. neg+unknown) | 1.185(0.395-3.559) | 0.762 | - | - |
| HSPB1 expression (High vs. Low) | 7.042(2.556-19.398) | **<0.001** | 5.520(1.984-15.355) | **0.001** |

Abbreviation: 95% CI=95% confidence interval; HR=hazard ratio; LN=lymph nodes; ER=estrogen receptor; PR=progesterone receptor; HER-2: human epidermal growth factor receptor-2; p value < 0.05 marked in bold font to show statistical significant.

**Table 3.** Univariate and multivariate Cox regression analyses for DFS of 131 breast cancer patients.

|  | Univariate analysis |  | Multivariate analysis |  |
| --- | --- | --- | --- | --- |
|  | HR(95%CI) | p value | HR(95%CI) | p value |
| Age($\geq 59$ vs. $< 59$) | 2.412(1.169-4.975) | **0.017** | 2.171(1.038-4.540) | **0.039** |
| Histologic grade |  |  |  |  |
| G2 vs. G1 | 9683.222<br>(0.000-1.018E+69) | 0.904 | - | - |
| G3 vs. G1 | 9420.873(0.000-9.911E+68) | 0.905 | - | - |
| Unknown vs. G1 | 5068.560(0.000-5.361E+68) | 0.911 | - | - |
| Tumor size($> T1$ vs. $\leq T1$+unknown) | 1.638(0.788-3.407) | 0.187 | - | - |
| LN metastasis |  |  |  |  |
| 1-3 vs. 0 | 2.956(1.162-7.520) | **0.023** | 3.677(1.348-10.033) | **0.011** |
| $> 3$ vs. 0 | 10.279(4.180-25.280) | **<0.001** | 7.422(2.808-19.617) | **<0.001** |
| Distant metastasis (Yes vs. No+unknown) | 6.412(3.105-13.241) | **<0.001** | 2.726(1.284-5.789) | **0.009** |
| ER status(pos vs. neg) | 1.531(0.683-3.432) | 0.301 | - | - |
| PR status(pos vs. neg) | 1.396(0.625-3.120) | 0.415 | - | - |
| HER-2 status(pos vs. neg+unknown) | 2.242(0.298-16.885) | 0.433 | - | - |
| Ki67 expression(pos vs. neg+unknown) | 0.932(0.398-2.183) | 0.871 | - | - |
| HSPB1 expression (High vs. Low) | 5.753(2.631-12.579) | **<0.001** | 4.279(1.886-9.709) | **0.001** |

Abbreviation: 95% CI=95% confidence interval; HR=hazard ratio; LN=lymph nodes; ER=estrogen receptor; PR=progesterone receptor; HER-2: human epidermal growth factor receptor-2; p value $< 0.05$ marked in bold font to show statistical significant.

### HSPB1 overexpression facilitates breast cancer cell proliferation, migration, and invasion in vitro

To investigate whether HSPB1 can alter breast cancer cell tumor biology, a series of in vitro experiments were performed in MDA-MB-231 and MDA-MB-468 cell lines overexpressing HSPB1 or with HSPB1 knockdown. Overexpression efficiency was confirmed by qRT-PCR and western blot (Figure 2A). The MTT and colony formation assays indicated that HSPB1 overexpression promoted the breast cancer cell proliferation (Figure 2B-C). Consequently, HSPB1 overexpression led to increased DNA synthesis activities as determined by Edu assays (Figure 2D). As shown in the wound healing assay and transwell assay, HSPB1 overexpression markedly promoted the migration and invasion abilities of both MDA-MB-231 and MDA-MB-468 cells (Figure 2E-F). The epithelial–mesenchymal transition (EMT) is the major mechanism of migration and invasion of cancer cells. Therefore, we further assessed the role of HSPB1 on the expression of EMT markers. Western blot assay (Figure 2G) revealed that HSPB1 overexpression led to decreased expression of epithelial markers (E-cadherin) and increased expression of mesenchymal markers (Fibronectin, N-cadherin, Vimentin), highlighting the significant effect of HSPB1 on regulating EMT in breast cancer cells. Furthermore, HSPB1 knockdown inhibited cell growth, migration, invasion, and EMT in breast cancer cells (Figure 2-figure supplement 1). Collectively, these results indicated that HSPB1 acted as a tumor promoter in breast cancer cells.

**Figure 2.**
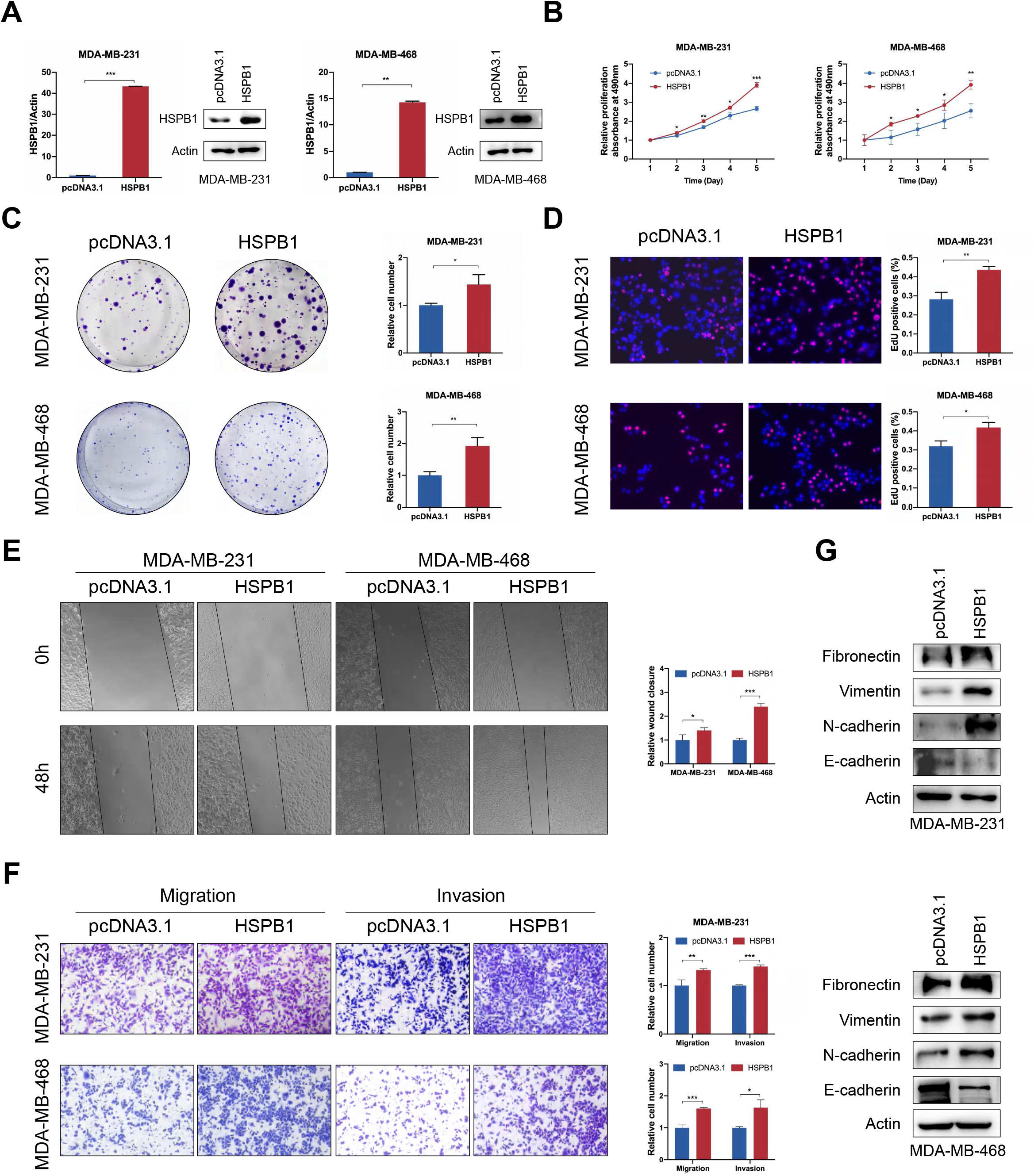
HSPB1 overexpression promoted breast cancer growth, migration, and invasion in vitro. (A) HSPB1 overexpression efficiency was confirmed by qRT-PCR and western blot in breast cancer cells. (B-D) MTT (B), colony formation (C), and EdU (D) assays were performed to evaluate the effect of HSPB1 overexpression on cell proliferative ability. (E) Wound healing assay was used to evaluate the effect of HSPB1 overexpression on the migration ability of breast cancer cells. (F) The migratory and invasive abilities of HSPB1 overexpressing breast cancer cells were assessed by transwell assay. (G) Western blot showed the effect of HSPB1 on the expression of EMT-related proteins. (* P < 0.05, ** P < 0.01, *** P < 0.001) **Figure 2-source data 1** Original western blots for Figure 2A. **Figure 2-source data 2** Original western blots for Figure 2G.

### HSPB1 overexpression suppresses ferroptosis to promote doxorubicin resistance of breast cancer cells in vitro

Elevated mRNA and protein expression levels of HSPB1 were identified in doxorubicin-resistant breast cancer cells compared with parental cells (Figure 3A-B), indicating its promoting role in doxorubicin resistance. Moreover, doxorubicin treatment led to increased HSPB1 expression in a dose-dependent and time-dependent manner (Figure 3B, Figure 3-figure supplement 1A-B). We also analyzed the expression level of HSPB1 in breast cancer tissues with different response to chemotherapy using several public breast cancer datasets, and the results showed that HSPB1 mRNA is upregulated in the majority of chemoresistant breast cancer tissues (Figure 3-figure supplement 1C). These results highlighted the significant association between HSPB1 expression and doxorubicin resistance. Therefore, we further evaluated the effect of HSPB1 on the breast cancer cell resistance to doxorubicin. Significantly, the IC_50_ value of doxorubicin was significantly higher in HSPB1 overexpressing breast cancer cells compared to that in control cells (Figure 3C). Consistently, breast cancer cells transfected with si-HSPB1 showed attenuated resistance to doxorubicin compared to cells transfected with si-NC (Figure 3D, Figure 3-figure supplement 2A). Moreover, HSPB1 knockdown could lead to decreased cell proliferation, migration and invasion of doxorubicin-resistant cells (Figure 3-figure supplement 2B-G). These results revealed the essential role of HSPB1 in doxorubicin resistance of breast cancer.

**Figure 3.**
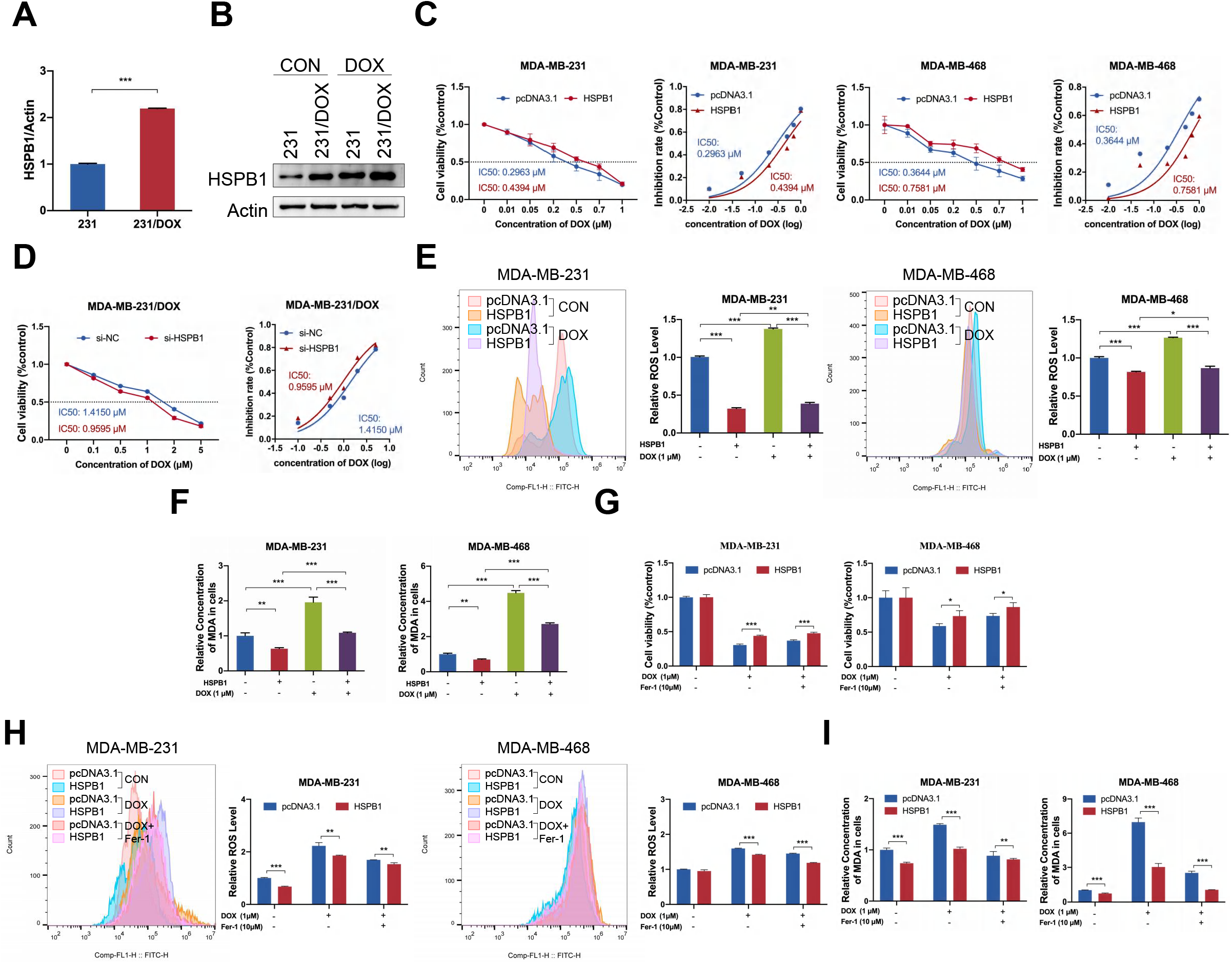
HSPB1 overexpression suppressed doxorubicin-induced ferroptosis in breast cancer cells. (A-B) The RNA (A) and protein (B) expression of HSPB1 was elevated in doxorubicin-resistant breast cancer cells. (C) Cell viability of MDA-MB-231 and MDA-MB-468 cells with or without HSPB1 overexpression was analyzed by MTT assay 48 h after treatment with different concentrations of doxorubicin. (D) Cell viability of MDA-MB-231/DOX cells with or without HSPB1 knockdown was analyzed by MTT assay 48 h after treatment with different concentrations of doxorubicin. (E-F) ROS levels (E) and cellular MDA levels (F) were determined in MDA-MB-231 and MDA-MB-468 cells with or without HSPB1 overexpression 48 h after treatment with 1 μM doxorubicin. (G) Cell viability was examined in MDA-MB-231 and MDA-MB-468 cells with or without HSPB1 overexpression 48 h after treatment with doxorubicin (1 μM) plus either DMSO or Ferrostatin-1 (Fer-1, 10 μM). (H-I) ROS levels (H) and cellular MDA levels (I) were determined in MDA-MB-231 and MDA-MB-468 cells with or without HSPB1 overexpression 48 h after treatment with 1 μM doxorubicin plus either DMSO or 10 μM Fer-1. (* P < 0.05, ** P < 0.01, *** P < 0.001) **Figure 3-source data 1** Original western blots for Figure 3B.

Recently, HSPB1 is identified as a novel negative regulator of ferroptosis in several human cancer cells (22). such as Hela, U2OS, and LNCap cells. Our results indicated that erastin (an inducer of ferroptosis) treatment led to increased mRNA and protein level of HSPB1 in a dose-dependent manner in breast cancer cells (Figure 3-figure supplement 3A), which is consistent with the observations in other cancer cells (22). HSPB1 overexpression inhibited while HSPB1 knockdown increased erastin-induced intracellular concentrations of lipid ROS in breast cancer cells (Figure 3-figure supplement 3B-C), which is one of the significant signatures of ferroptosis. Furthermore, we treated breast cancer cells with ferrostatin-1 (Fer-1, an inhibitor of ferroptosis). The results indicated that HSPB1 overexpression decreased erastin-induced growth inhibition and ROS production in breast cancer cells, and ferrostatin-1 treatment showed synergistic effect with HSPB1 overexpression (Figure 3-figure supplement 3D-E). Consistently, HSPB1 knockdown led to opposite results, and treatment with Fer-1 attenuated the effect of erastin in HSPB1 knockdown cancer cells (Figure 3-figure supplement 3D-E). The results observed in MDA-MB-231/DOX cells further confirmed the above findings (Figure 3-figure supplement 3F-H). Thus, these observations indicated significant role of HSPB1 in the regulation of ferroptosis in breast cancer cells.

Previous study has reported the association between doxorubicin and ferroptosis (23), however, the detailed anti-tumor effect of doxorubicin-induced ferroptosis is not fully elucidated. Therefore, we further investigated whether HSPB1 could regulate the anti-cancer activity of doxorubicin through modulating ferroptosis in breast cancer cells. Indeed, doxorubicin treatment led to increased level of ROS and malondialdehyde (MDA, the metabolite of lipid peroxidation), two surrogate markers for ferroptosis, indicating that doxorubicin could induce ferroptosis in breast cancer cells (Figure 3E-F, Figure 3-figure supplement 4A-B). Significantly, the effect caused by doxorubicin treatment was attenuated by HSPB1 overexpression and enhanced by HSPB1 knockdown (Figure 3E-F, Figure 3-figure supplement 4A-B). Furthermore, Fer-1 was added in the HSPB1-overexpressing or HSPB1-knockdown cells with the presence of doxorubicin. The results showed that Fer-1 treatment could attenuate the effect of doxorubicin-induced ferroptosis (Figure 3G-I, Figure 3-figure supplement 4C-E), as indicated by the recovery of cell viability and the reduction of cellular ROS and MDA levels. Moreover, Fer-1 exhibited a synergistical promoting effect with HSPB1 overexpression and had an antagonistic effect with HSPB1 knockdown (Figure 3G-I, Figure 3-figure supplement 4C-E). As expected, the similar results of HSPB1 knockdown were found in MDA-MB-231/DOX cells, which was reflected in the inhibited cell viability and upregulated ROS and MDA levels (Figure 3-figure supplement 4F-H). We also investigated the role	of HSPB1 in paclitaxel (PTX)-induced ferroptosis, which is another frequently-used chemotherapeutics for breast cancer patients (24). The results indicated that HSPB1 could promote the resistance to PTX through inhibiting PTX-induced ferroptosis, as determined by the cell viability and ROS production under different condition (Figure 3-figure supplement 5). Altogether, these results strengthened the effect of HSPB1 on doxorubicin resistance through regulating ferroptotic cell death in breast cancer cells.

### HSPB1 regulates the activation of NF-κB signaling

In order to elucidate the underlying mechanisms of HSPB1-mediated resistance to doxorubicin-induced ferroptosis in breast cancer cells, we investigated the effect of HSPB1 on the activation of NF-κB signaling, which plays significant role in the progression of cancer (25). HSPB1 overexpression led to decreased expression of IKβ-α (a negative regulator of NF-κB) and increased expression of NF-κB, p-IKβ-α, and downstream molecules of NF-κB (Twist1, IL6, and Survivin) in breast cancer cells (Figure 4A), indicating the potential role of HSPB1 in regulating NF-κB activity. Using NF-κB reporter assay, we found that doxorubicin treatment led to upregulated NF-κB transactivation activity, and HSPB1 overexpression further enhanced the effect induced by doxorubicin (Figure 4B). Consistently, the protein levels of p-IKβ-α, NF-κB, and its target genes were increased and the protein expression of IKβ-α was decreased followed doxorubicin treatment, and HSPB1 overexpression led to increased effect of doxorubicin (Figure 4C). The qRT-PCR and IF assays also indicated the similar expression changes of Survivin and IL6 in breast cancer cells (Figure 4D-E). Moreover, HSPB1 knockdown could diminish the effect of doxorubicin treatment on the activation of NF-κB signaling (Figure 3-figure supplement 1). These data indicated that HSPB1 was involved in the activation of NF-κB signaling.

**Figure 4.**
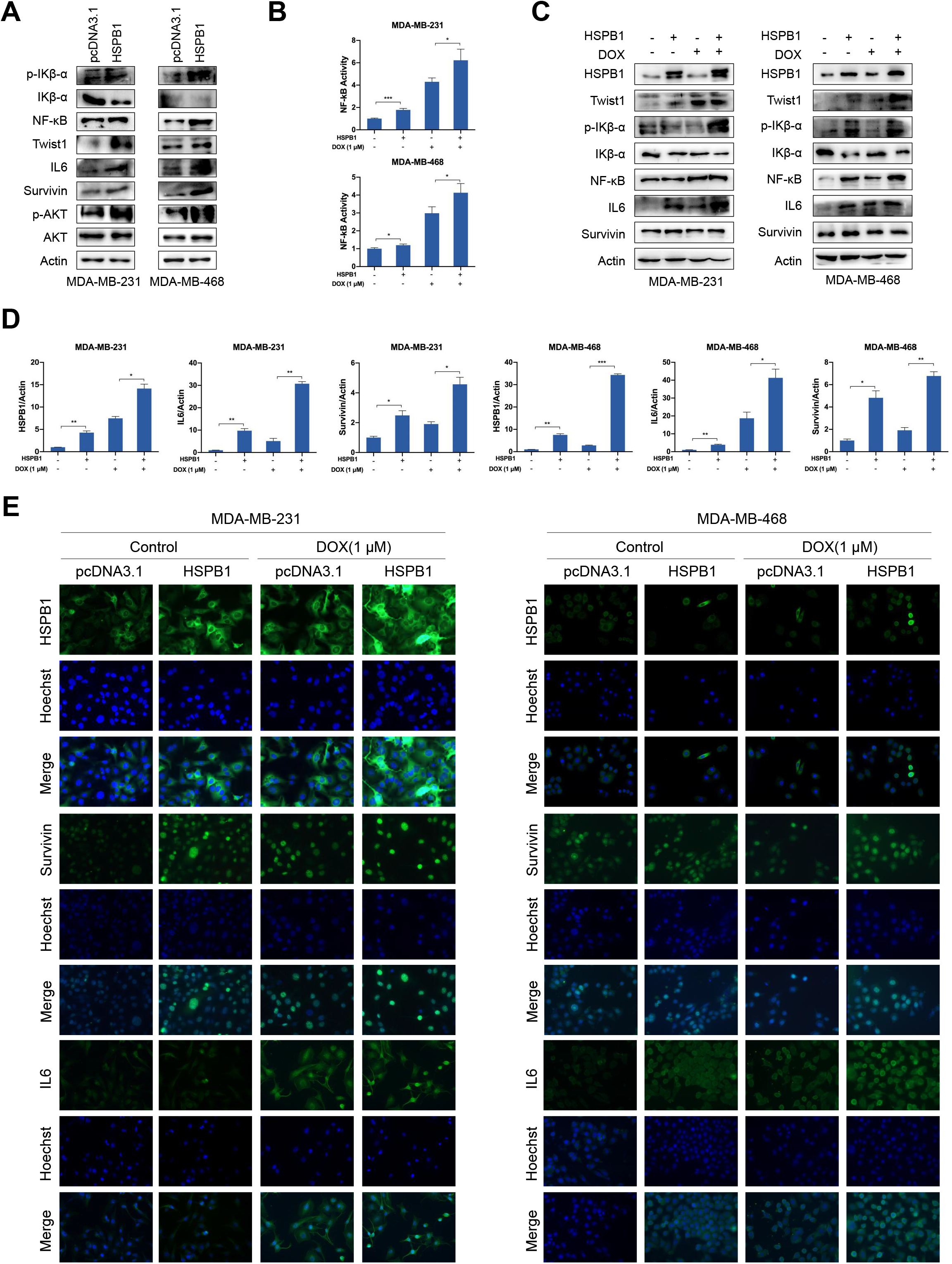
HSPB1 overexpression enhanced activation of NF-κB signaling. (A) Western blot was performed using cell lysates of MDA-MB-231 and MDA-MB-468 cells with or without HSPB1 overexpression. (B-D) NF-κB transcriptional activity was determined by NF-κB activation reporter assay (B), western blot (C), and qRT-PCR (D). (E) After 24h of treatment with 1 μM doxorubicin, immunofluorescence was performed to detect the expression of HSPB1, Survivin, and IL6 in MDA-MB-231 and MDA-MB-468 cells with or without HSPB1 overexpression. (* P < 0.05, ** P < 0.01, *** P < 0.001) **Figure 4-source data 1** Original western blots for Figure 4A. **Figure 4-source data 2** Original western blots for Figure 4C.

### HSPB1 promotes the ubiquitination-mediated degradation of Ikβ-α, leading to increased nuclear translocation of NF-κB in breast cancer cells

The IKβ-α could bind with NF-κB to keep it in the cytoplasm, preventing nuclear translocation of NF-κB. There is evidence showing that HSPB1 could enhance the ubiquitination-mediated degradation of Ikβ-α through binding with phosphorylated Ikβ-α, leading to release of NF-κB from the cytoplasmic NF-κB/Ikβ-α complex and enhanced NF-κB activity (26). We also revealed that doxorubicin treatment led to increased nuclear translocation of NF-κB, which was inhibited by HSPB1 knockdown (Figure 5A-B). Moreover, the co-IP assay demonstrated that doxorubicin treatment could attenuate the binding between Ikβ-α and NF-κB, while HSPB1 knockdown could restore the inhibited interaction of Ikβ-α with NF-κB (Figure 5C). In addition, the results also indicated an increase in the binding activity between HSPB1 and Ikβ-α induced by doxorubicin treatment (Figure 5D). Given our above-mentioned results (Figure 4A, Figure 3-figure supplement 1A), which revealed obvious effect of HSPB1 on the expression of Ikβ-α in breast cancer cells, we further examine the role of HSPB1 in regulating protein stability of Ikβ-α. 10 μM MG132 (a proteasome inhibitor) or 20 μM chloroquine (CQ, a lysosome inhibitor) was used to treat breast cancer cells to investigate the possibility of its proteasomal or lysosomal degradation. The results showed that overexpression of HSPB1 led to decreased expression of Ikβ-α in control and CQ group (Figure 5E). MG132 treatment led to upregulated expression of Ikβ-α compared to other two groups, however, the expression of Ikβ-α was less affected by HSPB1 overexpression in MG132 group (Figure 5E). Next, breast cancer cells transfected with empty vector or with HSPB1 overexpressing vector were treated with 100 μg/ml cycloheximide (CHX, a protein synthesis inhibitor). The results showed that overexpression of HSPB1 markedly decreased the half-life of Ikβ-α from 6.39 h (CHX + Vector) to 5.72h (CHX + HSPB1) in MDA-MB-231 cells and from 19.34 (CHX + Vector) to 8.08 (CHX + HSPB1) in MDA-MB-468 cells (Figure 5F), suggesting that HSPB1 could decrease the protein stability of Ikβ-α. We further examined whether HSPB1 regulated the ubiquitination of Ikβ-α in breast cancer cells. The results indicated that HSPB1 knockdown decreased the ubiquitinated levels of Ikβ-α compared to the control group (Figure 5G). These findings indicated that HSPB1 regulated the activity of NF-κB through contributing to ubiquitination-mediated degradation of Ikβ-α in breast cancer cells.

**Figure 5.**
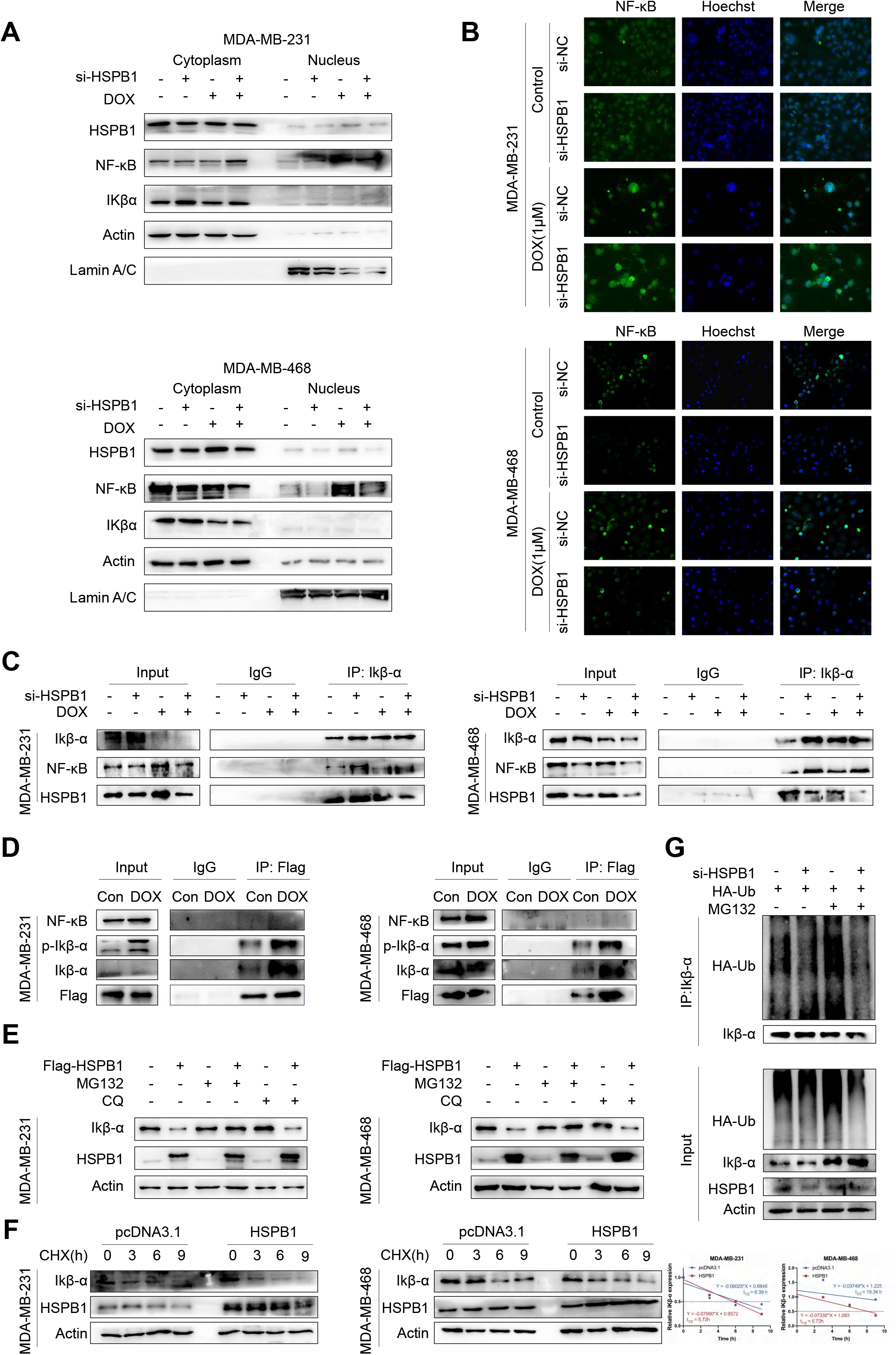
HSPB1 was involved in nuclear translocation of NF-κB through promoting ubiquitination-mediated degradation of Ikβ-α. (A) The protein levels in nuclear and cytoplasmic lysates were detected by western blot. Actin was used as an internal control for cytoplasmic lysates, and Lamin A/C was used as a loading control for nuclear lysates. (B) After 24h of treatment with 1 μM doxorubicin, MDA-MB-231 and MDA-MB-468 cells with or without HSPB1 knockdown were stained with anti-NF-κB antibody to investigate subcellular localization and expression levels. (C) Coimmunoprecipitation (Co-IP) analysis of Ikβ-α in cells treated with 1 μM doxorubicin for 24 h were evaluated for the presence of Ikβ-α, NF-κB, and HSPB1. (D) The interaction between HSPB1 and Ikβ-α or NF-κB in MDA-MB-231 and MDA-MB-468 cells after treatment with 1 μM doxorubicin was determined by Co-IP assay. (E) MDA-MB-231 and MDA-MB-468 cells were treated with MG132 or CQ, and western blot was used to detect the effect of HSPB1 on the expression of Ikβ-α. (F) MDA-MB-231 and MDA-MB-468 cells were transfected with HSPB1-overexpressing vectors or control vectors. After treatment with 100 μg/ml cycloheximide (CHX) for the indicated time, cells were collected for WB analysis. ImageJ software was used to quantify band intensity. (G) HEK293T cells with or without HSPB1 knockdown were simultaneously transfected with HA-Ub expression plasmids. Then, cells were treated with or without 10 μM MG132 for 6 h and collected for immunoprecipitation with anti- Ikβ-α antibody and ubiquitination detection. **Figure 5-source data 1** Original western blots for Figure 5A. **Figure 5-source data 2** Original western blots for Figure 5C. **Figure 5-source data 3** Original western blots for Figure 5D. **Figure 5-source data 4** Original western blots for Figure 5E. **Figure 5-source data 5** Original western blots for Figure 5F. **Figure 5-source data 6** Original western blots for Figure 5G.

### Restoring NF-κB activity attenuates the suppressive effect of HSPB1 knockdown in breast cancer cells

We further examined whether additional activation of NF-κB could diminish the inhibitory effect caused by HSPB1 knockdown. The increased expression of Ikβ-α and decreased expression of NF-κB caused by HSPB1 knockdown could be restored by knockdown of Ikβ-α (Figure 6A). Moreover, Ikβ-α knockdown could attenuate the inhibited effect of HSPB1 knockdown on cell proliferation, migration, and invasion (Figure 6B-E, Figure 6-figure supplement 1A). The NF-κB activity could be restored in HSPB1 knockdown cells by knockdown of Ikβ-α with or without doxorubicin treatment (Figure 6F-I), as determined by NF-κB reporter assay and detection of the expression of NF-κB target genes. In addition, the inhibited cell viability and enhanced ROS production induced by doxorubicin or erastin in HSPB1 knockdown cells could be restored by simultaneous knockdown of Ikβ-α (Figure 6J-K, Figure 6-figure supplement 1B). These results suggest that HSPB1 regulated the biological behaviors and doxorubicin-induced ferroptosis through NF-κB activity in breast cancer cells.

**Figure 6.**
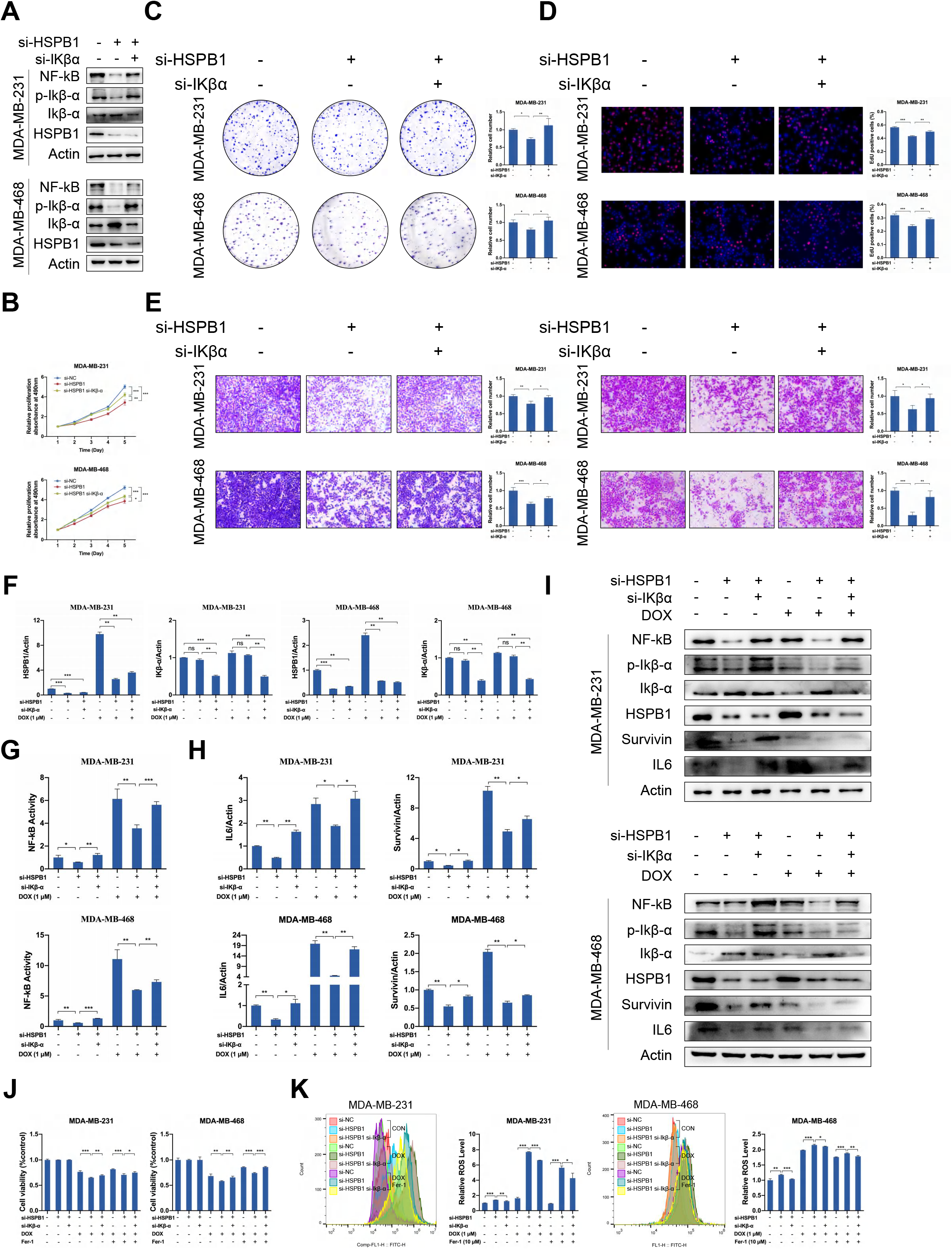
Ikβ-α knockdown partially restored the suppressive effect of HSPB1 knockdown in breast cancer cells. MDA-MB-231 and MDA-MB-468 cells were transfected with negative control siRNA (Ctrl-siRNA), ctrl-siRNA + si-HSPB1, or si-HSPB1 + si-Ikβ-α for 48 h. (A) The expression levels of HSPB1, Ikβ-α, p-Ikβ-α, and NF-κB were determined by Western blot. (B-D) The MTT assay (B), colony formation assay (C), and Edu assay (D) were used to detect cell proliferative ability. (E) Transwell assay was performed to determined cell migration and invasion. (F) The transfected MDA-MB-231 and MDA-MB-468 cells were treated with or without 1 μM doxorubicin. The HSPB1 and Ikβ-α expression was detected by qRT-PCR in indicated cells. (G) NF-κB transactivation was detected by NF-κB activation reporter assay. (H) The RNA expression levels of Survivin and IL6 were detected by qRT-PCR. (I) The protein expression levels of HSPB1, Ikβ-α, p-Ikβ-α, NF-κB, Survivin and IL6 were determined by Western blot. (J) The MTT assay was used to analyze the viability of transfected cells following doxorubicin plus either DMSO or Fer-1 treatment. (K) Cellular ROS levels were detected in transfected cells after doxorubicin plus either DMSO or Fer-1 treatment. (* P < 0.05, ** P < 0.01, *** P < 0.001) **Figure 6-source data 1** Original western blots for Figure 6A. **Figure 6-source data 2** Original western blots for Figure 6I.

### HSPB1 enhances malignant behaviors of breast cancer cells by regulating IL6 expression

IL6 belongs to the family of IL (interleukin)–6-type cytokines, which could be secreted by myriads of cells (27, 28), including monocytes, fibroblasts, endothelial cells, keratinocytes, and cancer cells. Previous studies reported that IL6 showed significant role in regulating various cellular functions (29, 30), such as cell proliferation, metastasis, vascular permeability, metabolism, and infiltration of immune cells. Given the obvious effect of HSPB1 on the expression of IL6 in breast cancer cells detected by our above results, we further investigate whether the secretion of IL6 could be influenced by HSPB1. The ELISA assay indicated that HSPB1 knockdown led to decreased IL6 secretion while HSPB1 overexpression promoted the secretion of IL6 (Figure 7A). Next, we evaluated the influence of secreted IL6 on the malignant behaviors of breast cancer. The supernatants from HSPB1 overexpressing cells led to increased migration ability of breast cancer cells and tube formation of HUVECs, while supplement of IL6 neutralizing antibodies could partly attenuate the promoted effect (Figure 7B).

**Figure 7.**
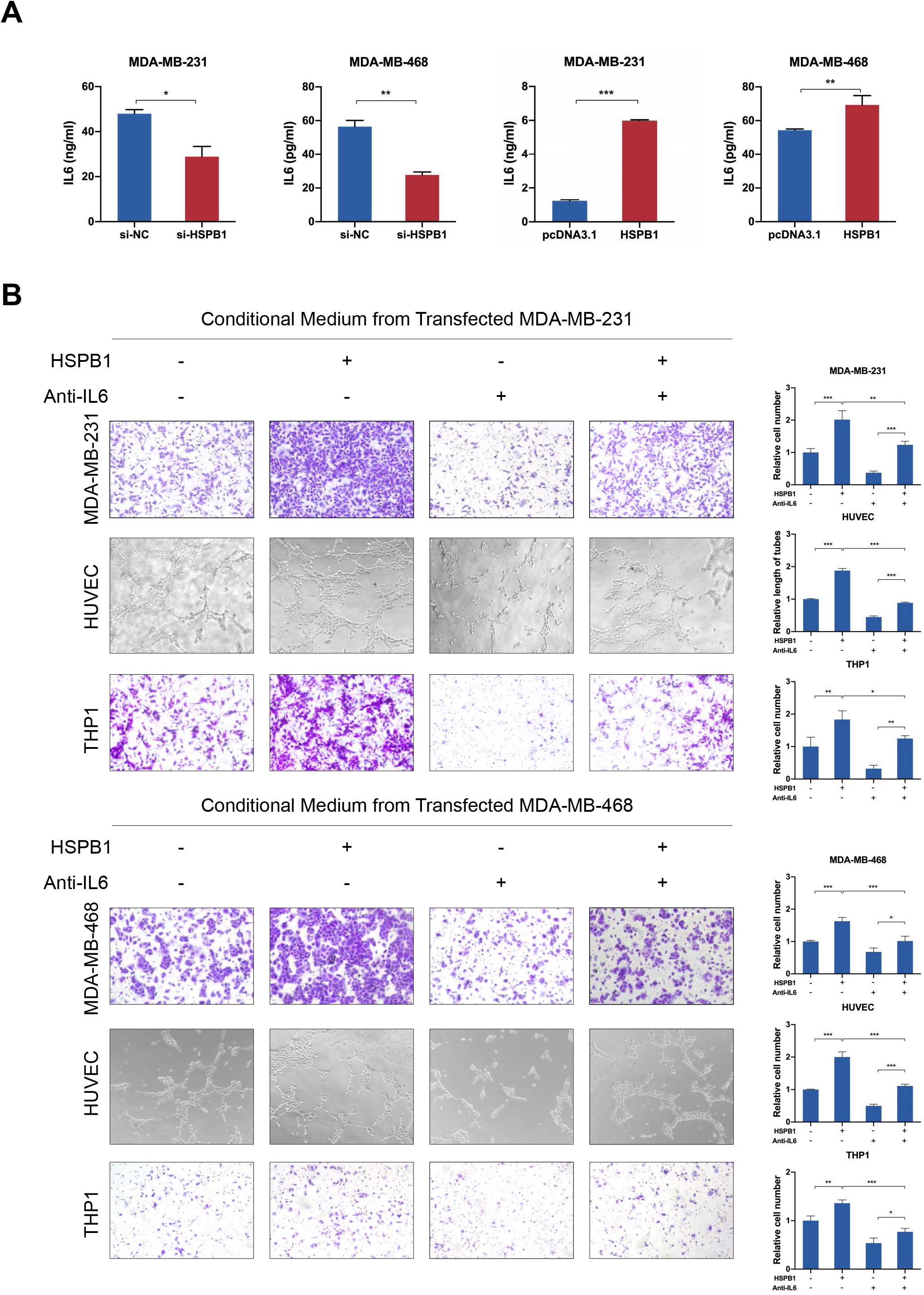
HSPB1 promoted breast cancer progression through regulating IL6. (A) The ELISA assays were conducted to detect the expression levels of IL6 in transfected cells. (B) IL6 neutralizing antibody treatment (2.5 μg/ml) efficiently reversed the promoting effects of conditioned medium from HSPB1-overexpressing MDA-MB-231 or MDA-MB-468 cells on the breast cancer cell migration, tube formation, and THP1 migration. (* P < 0.05, ** P < 0.01, *** P < 0.001)

As the major component of tumor microenvironment, tumor-associated macrophages play an important role in the whole process from tumor occurrence to metastasis (31). We used CIBERSORT, xCell, and quanTIseq algorithms to evaluate the levels of immune cell infiltration in tumor microenvironment. The results of Spearman correlation analysis showed that the expression level of HSPB1 was positively correlated with the infiltration level of M2 macrophages and the ratio of M2 to M1, while negatively correlated with the infiltration level of M1 macrophages (Figure 7-figure supplement 1A). It could be speculated that HSPB1 might play an important role in the progression of breast cancer by promoting the infiltration of M2 macrophages. Previous studies reported that tumor cells could produce numerous chemokines to attract macrophages, which further dictate the fate of tumor developing and progression through secreting an assorted array of cytokines. It is well-established that IL6 is an essential chemokine for the recruitment of macrophages in cancers (30, 32), such as cervical cancer and prostate cancer. Accordingly, we further evaluated whether HSPB1 could modulate the infiltration of macrophage in breast cancer. The conditioned medium from HSPB1 knockdown cells led to decreased effect on the migration ability of macrophages and attenuated chemotaxis to macrophages, while the conditioned medium from HSPB1 overexpressing cells exhibited the opposite effect (Figure 7-figure supplement 1B-E). To confirm that the enhanced behaviors of macrophages induced by HSPB1 was mediated by IL6, anti-IL6-neutralizing antibodies were used. The results showed that the addition of IL6 neutralizing antibodies could abrogate the enhanced effect of supernatants from HSPB1 overexpressing cells on THP1 migration (Figure 7B), suggesting the involvement of HSPB1-IL6 pathway in the regulation of accumulation of macrophages in breast cancer. Collectively, these observations indicated that HSPB1 could aggravate progression of breast cancer through promoting IL6 secretion.

### HSPB1 promotes tumor growth, doxorubicin resistance, and metastasis of breast cancer in vivo

We further evaluated the function of HSPB1 in breast cancer in vivo using a nude mouse xenograft model. The stable HSPB1-overexpressing or control MDA-MB-231 cells were injected into the flanks of BALB/c nude mice. When the tumor volume reached 50 mm^3^, PBS or doxorubicin was injected intravenously. As shown in Figure 8a-c, HSPB1 overexpression led to increased tumor volume and tumor weight. Moreover, the tumors in the doxorubicin-injected mice were smaller and lighter than those in PBS-treated mice (Figure 8A-C), indicating the suppressive effect of doxorubicin on the growth of breast cancer in vivo. Significantly, the tumors in the doxorubicin-injected mice bearing HSPB1-overexpressing cells were significantly larger and heavier compared to those in doxorubicin-injected mice implanted with control cells (Figure 8A-C), suggesting that overexpression of HSPB1 attenuated the anti-tumor effect of doxorubicin. HE staining was used to evaluated the morphology of the tumors (Figure 8D). The IHC analysis indicated that the expression changes of IKβ-α, NF-κB and its target genes in the tumor tissues were consistent with the findings in vitro (Figure 8D). Moreover, doxorubicin decreased the protein expression of Ki-67, which was enhanced by HSPB1 overexpression (Figure 8D). These results demonstrated that HSPB1 increased the resistance of breast cancer to doxorubicin-induced ferroptosis via NF-κB signaling in vivo.

**Figure 8.**
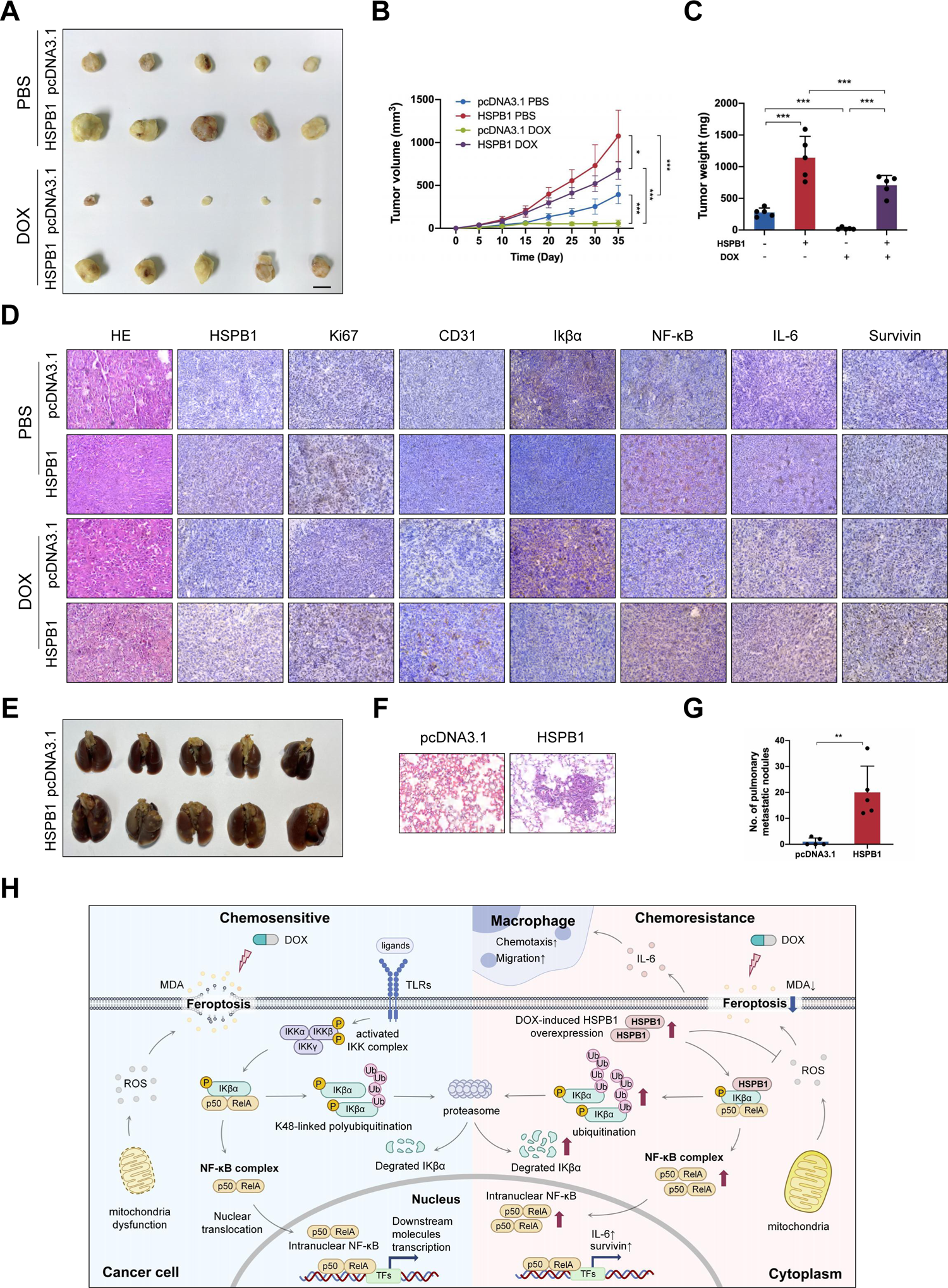
HSPB1 promoted chemoresistance and metastasis of breast cancer. (A) Control or HSPB1-overexpressing MDA-MB-231 cells were transplanted into the flanks of nude mice, followed by doxorubicin or PBS treatment. Images of tumors harvested. (B) The tumor volumes were measured every five days. (C) Tumor weights were recorded after sacrifice of the mice. (D) H&E staining showed the tissue morphology of transplanted tumors. Representative pictures of immunohistochemical staining of HSPB1, Ki67, CD31, Ikβα, NF-κB, IL-6, and Survivin in the tumor tissues. (E) Pictures of lung metastatic nodules. (F) H&E staining showed the tissue morphology of lungs isolated from indicated mice. (G) The statistical graph of lung metastatic nodules. (H) The mechanistic schematic model of the role of HSPB1 in breast cancer. (* P < 0.05, ** P < 0.01, *** P < 0.001)

In addition, we also investigated the effect of HSPB1 on breast cancer metastasis in vivo using a pulmonary metastasis model. The stable HSPB1-overexpressing or control MDA-MB-231 cells were injected into BALB/c nude mice via the tail vein. The lung metastatic burden was significantly higher in HSPB1-overexpressing mice, in which the number of metastatic mice and lung metastatic foci were increased compared to those in control group (Figure 8E-G). Hematoxylin and eosin (H&E) staining was used to pathologically confirm the metastatic nodule in the dissected lungs, and overexpression of HSPB1 significantly increased the size and number of lung metastatic nodules (Figure 8E-G). Collectively, these results suggested that HSPB1 facilitated tumor growth, doxorubicin resistance, and metastasis in vivo (Figure 8H).

## Discussion

Due to the lack of ER, PR, and HER2 receptors, there is no ideal target available for effective therapy against TNBC (33). Chemotherapy remains the first-line treatment for most TNBC patients (3, 34). However, TNBC is an aggressive phenotype and often refractory to chemotherapy, leading to disease progression, relapse and metastasis, which is the reason for more than 90% death of TNBC patients (35, 36). Therefore, finding novel targets that could clarify the underlying mechanisms and be exploited to overcome chemoresistance are critical for improving the prognosis of breast cancer patients. In the present study, we substantiated that HSPB1 could prevent breast cancer cells from chemotherapy-induced ferroptosis, and exhibited significant role in mediating progression and chemoresistance of breast cancer.

HSPB1 is a member of the small heat shock proteins, which could be highly inducible under stressful conditions (37). Previous studies have shown that HSPB1 exerted a tumor-promoting effect in various cancers through its proliferative and anti-apoptotic functions, such as esophageal squamous cell (37), prostate cancer (38), carcinoma squamous cell carcinoma of tongue (39). However, the effect of HSPB1 on the progression and chemoresistance and the underlying mechanism in breast cancer has not yet been fully explored. In the study, we identified upregulated HSPB1 expression in breast cancer tissues based on the results obtained from several public datasets, which was further confirmed using tissues from our cohort. Moreover, high expression of HSPB1 was associated with worse prognosis of breast cancer patients. Subsequently, our results revealed that HSPB1 could promote breast cancer proliferation, migration, invasion, and doxorubicin resistance.

Doxorubicin (DOX) is a cytotoxic anthracycline antibiotic, widely used for treating a variety of cancers, which is considered as the foundation of chemotherapy for breast cancer (40). Accumulating studies have shown that doxorubicin promoted cancer cell death through inhibiting DNA replication and generating H_2_O_2_, which is the Fenton reaction substrate leading to increased ROS production (41, 42). Thus, doxorubicin might kill cancer cells by indirectly inducing ferroptosis. Previous studies reported that HSPB1 might be a negative regulator of ferroptosis (22), which was induced upregulation following erastin treatment in several cancer cells, such as Hela cells, U2OS and LNCap. Here, we observed similar functions of doxorubicin in breast cancer cells. Doxorubicin treatment substantially increased the intracellular ROS and MDA levels in breast cancer cells, which are the critical executors of ferroptosis (14). In consistent with the previous study, we identified the induced expression of HSPB1 after erastin treatment (22). In addition, our results also revealed the doxorubicin-mediated upregulation of HSPB1 in breast cancer, indicating that HSPB1 might be involved in the regulation of doxorubicin-induced ferroptosis. In fact, HSPB1 overexpression led to attenuated doxorubicin-induced ferroptosis, as determined by the reduction of cellular concentrations of lipid ROS and MDA, and treatment with Fer-1 could further enhance the effect caused by HSPB1 overexpression. These results suggested that HSPB1 overexpression inhibited ferroptosis and ultimately led to a decrease of the doxorubicin sensitivity of breast cancer cells. We further constructed the doxorubicin-resistant cells (MDA-MB-231/DOX), and significantly upregulated expression of HSPB1 was identified in the chemoresistant cells compared to the corresponding parental cells. HSPB1 knockdown could render chemoresistant cells regain sensitivity to doxorubicin treatment through promoting doxorubicin-induced ferroptosis. Similar findings have been observed following paclitaxel treatment, another wildly-used anti-cancer drug, indicating the significant role of HSPB1 in chemotherapy-induced ferroptosis.

NF-κB pathway is essential for governing inflammatory response in cancers, and is implicated as a hallmark of cancer progression and a potential therapeutic target (43). To gain insight into the intricate molecular mechanism by which HSPB1 regulated doxorubicin-induced ferroptosis, we first explored whether HSPB1 was able to regulate NF-κB pathway to exhibit tumor-promoting role in breast cancer. Here, we found that HSPB1 could promote the nuclear translocation and activity of NF-κB in breast cancer cells, as determined by the elevated expression of its target genes. Previous study reported that ubiquitin‐mediated degradation of NF‐κB inhibitor proteins, known as IκBs, was the major cause of activation of NF-κB signaling (44). Among the IκB family, IκB-α is the most well‐studied member. HSPB1 was reported to promote the 26S proteasome-mediated degradation of ubiquitinated proteins, such as phosphorylated IKβ-α (26). Consistently, our results also demonstrated that HSPB1 could bind with IKβ-α and promote ubiquitination-mediated IKβ-α degradation, leading to enhanced nuclear translocation and activity of NF-κB in breast cancer cells. Moreover, doxorubicin treatment could further reinforce the mutual combination between HSPB1 and IKβ-α, indicating the underlying mechanism involved in HSPB1-mediated doxorubicin resistance. In addition, the inhibitory effect caused by HSPB1 knockdown could be reversed by knockdown of IKβ-α in breast cancer cells, which further demonstrated that HSPB1 regulated the aggressive behaviors of breast cancer cells through NF-κB activity.

NF-κB is a well-known transcriptional factor involved in the expression of several cytokines. In this study, high expression of IL6 was identified in HSPB1-overexpressing group, and the inhibited expression of IL6 caused by HSPB1 knockdown could be rescued by IKβ-α knockdown. IL6 could be secreted from various cells in cancer tissues, including myeloid cells, cancer-associated fibroblasts, and cancer cells (45–47). Our results revealed that HSPB1 overexpression promoted the secretion of IL6, whereas HSPB1 knockdown led to decreased IL6 secretion in breast cancer cells. Previous study reported that high serum IL6 concentration was associated with poor prognosis of breast cancer patients (48). Moreover, IL6 family members was reported to promote cell migration, angiogenesis, as wells as macrophage recruitment and infiltration (49, 50). Therefore, we further investigated the effect of HSPB1-induce IL6 in the progression and immune infiltration of breast cancer. The conditioned medium collected from HSPB1-overexpressing cells significantly promoted the migration of cancer cells and macrophages as well as angiogenesis of HUVECs, which could be abrogated by the supplement of IL6 neutralizing antibodies. IL6 has also been reported to induce macrophage polarization to the M2-phenotype in various cancers (51, 52), however, the association between HSPB1 and M2 polarization in breast cancer needed to be further elucidated. Overall, IL6 showed a significant role in the HSPB1-mediated malignant behaviors, and it is possible that HSPB1 is involved in the regulatory function of IL6 through modulating NF-κB activity in breast cancer.

In conclusion, our results uncovered that HSPB1 is a significant drug-resistant factor, which could inhibit chemotherapy-induced ferroptosis in breast cancer through promoting the activation of NF-κB signaling pathway. Our observations provide a foundational rationale that targeting HSPB1 and combinational induction of ferroptosis with anticancer drugs would be a potential therapeutic strategy to overcome chemoresistance in breast cancer.

## Materials and methods

### Public data access and analysis

The gene expression profiles and prognostic information of breast cancer patients were downloaded from TCGA (https://portal.gdc.cancer.gov/) and GEO (http://www.ncbi.nlm.nih.gov/gds/) databases. Limma package of R software was used for the analysis of differentially expressed mRNAs (DEmRNAs) between normal samples and normal samples. The criteria for selection of DEmRNAs was log_2_|fold change (FC)|≥ 1 and adjust P value < 0.05. Heatmap package of R software was used to draw heatmaps.

### Patients and samples

The breast cancer tissues and normal tissues were obtained from patients diagnosed with breast cancer undergoing surgery at the Qilu hospital of Shandong university from March 2007 to January 2017. Survival time was defined as the time from the date of operation until the date of last follow-up or death. This study was approved by the Ethical Committee of Qilu Hospital of Shandong University (KYLL-2016-255). The informed consents for participation in this study from all patients were obtained.

### Cell culture and transfection

Human breast cancer cell lines (MDA-MB-231 and MDA-MB-468), human umbilical vein endothelial cells (HUVECs), THP-1 cells, and HEK293T cells were purchased from American Type Culture Collection (ATCC, USA). The cells were authenticated by STR analysis (Suzhou, China) and tested for negative mycoplasma contamination using Mycoplasma Detection Kit (Sigma). MDA-MB-231, MDA-MB-468, HUVEC, HEK293T cell lines were cultured in DMEM (Macgene, China), and THP1 cells were maintained with RPMI-1640 medium. The culture medium was supplemented with 10% fetal bovine serum (FBS, Gibco, USA), 100 U/ml penicillin, and 100 μg/ml streptomycin. These cells were incubated in a humidified incubator containing 5% CO_2_ at 37 °C.

The full length of HSPB1 was cloned into pcDNA3.1 (Invitrogen, USA) to generate pcDNA3.1-HSPB1 constructs. After transfection with the overexpression vectors or empty vectors, cells were selected by G418 (1 mg/ml) for 2 months until the formation of visible clones. The survived cell clones were amplified and the overexpression efficiency of individual clones was confirmed using qRT-PCR assays. The satisfactory clones were identified as stably transfected cells and used for in vivo experiments. The siRNAs were purchased from GenePharma (Shanghai, China), and the target sequences for HSPB1, Ikβ-α, and Control are listed in Table S1. Lipofectamine 2000 (Invitrogen, USA) was used for cell transfection following the manufacturer’s instructions.

### Cell proliferation assays

Cell proliferation assays were performed using MTT assay. Briefly, 1,500 transfected cells were seeded into a 96‐well plate. Cell proliferation was assessed after indicated time. The cells were incubated for 4-6 h after the addition of 20 μl MTT (5 mg/ml) per well. Then, the culture supernatant was removed and 100 μl DMSO was added into each well. The absorbance was measured at 490 nm using a Microplate Reader (PerkinElmer, USA).

### IC_50_ detection and cytotoxic assay

The transfected cells were cultured in 96-well plates (3,000 cells/well). For IC_50_ detection, cells were grown in complete medium containing different concentrations of doxorubicin (MCE, USA) or paclitaxel (MCE, USA) for 48 h. The cell survival rate was further evaluated by MTT assay. For cytotoxic assay, cells were treated differently as indicated using doxorubicin, paclitaxel, or erastin (MCE, USA) with or without ferrostatin-1 (MCE, USA) for 48 h. The cell viability was assessed by MTT assay and presented as relative to the absorbance value of the control group.

### Colony formation assay

1,000 Transfected cells were seeded into 6 cm plates and cultured for over 2 weeks. Then, cells were washed with PBS, fixed with methanol for 15 min and subsequently stained with 0.2% crystal violet for 20 min at room temperature. Colonies were counted and photographed.

### EdU incorporation assay

The measurement of proliferative cells and nucleic acid were performed using EdU incorporation assay kit (RiboBio, China) according to the manufacturer’s instructions. Briefly, 2 × 10^4^ transfected cells were seeded into a 96-well plate per well for 24 h. After incubation with 50 μM EdU for another 2.5 h, cells were washed with PBS, fixed with 4% paraformaldehyde (PFA) for 30 min, and permeabilized with 0.5% Triton X-100 for 10 min. Subsequently, cells were stained with 1× Apollo Dye Solution for 30 min and 1× Hoechst for 30 min at room temperature in the dark. The images were obtained under a fluorescence microscope (ZEISS, Germany) and positive cells were counted.

### Wound healing assay

The transfected cells were seeded in a 24-well culture plate and cultured until reaching to 90% confluence. The cell monolayer was scratched using a sterile tip (10 μl), washed with PBS, and cultured with serum-free DMEM. Photographs were taken with a light microscope at the indicated time.

### Migration and invasion assays

For migration assays, 80,000 transfected cells resuspend with serum-free DMEM were seeded into the upper insert of transwell chamber (pore size 8 μm; Corning, China). 700 μl medium supplemented with 20% FBS were put into the lower chambers. For invasion assays, the transwell chambers were coated with Matrigel (Corning, China) solution. After incubation for 24-48h, the cells penetrated to the lower surface of the membrane were fixed with methanol for 15 min and then stained with 0.2% crystal violet for 20 min. Then the stained cells were photographed and counted.

### RNA preparation and qRT‐PCR

Total RNA was extracted from cells or frozen tissues using the TRIzol Reagent (Takara, Japan). The complementary DNA (cDNA) was synthesized using a PrimeScript reverse transcriptase reagent kit (Takara, Japan). The expression of mRNA was examined by real-time quantitative-PCR (qRT-PCR) using SYBR Green PCR Master Mix (Takara, Japan) and a LightCycler480 Detection System (Roche, Germany). The relative expression of indicated genes was analyzed by the 2^−ΔΔCt^ method. β-actin was used as normalization control. The primer sequences used for qRT-PCR are listed in Table S2.

### Western blot analysis

Total protein was extracted from cells or frozen tissues using the RIPA lysis buffer (Beyotime, China) with protease inhibitor (PMSF) and phosphatase inhibitor (NaF). The protein concentration was determined using a BCA protein assay kit (Beyotime, China). The protein samples were separated by 10% SDS-PAGE gel and then transferred to a 0.22 μm polyvinylidene difluoride (PVDF) membrane (Millipore, USA) in a wet electron transfer device. Then, the membrane was blocked using 5% skimmed milk in TBST for 1 h at room temperature and incubated with the primary antibody overnight at 4 °C. The next day, after incubation with the secondary antibody for 1 h at room temperature, an enhanced chemiluminescence (ECL) kit (Vazyme, China) was used to visualize the target protein. The antibodies are listed in Table S3.

### Immunofluorescence staining

5×10^5^ cells were seeded into 24-well culture plates for 24 h, and treated with the indicated drugs or left untreated for another 24 h. The cells were washed with PBS, fixed with 4% PFA for 30 min, and permeabilized with 0.5% Triton X-100 for 10 min. Subsequently, cells were blocked with 10% goat serum in PBS at room temperature for 1 h, and incubated with primary antibody at 4 °C overnight. After incubation with secondary antibody for 2 h at room temperature in the dark, the nuclei were stained by DAPI (Beyotime, China) at room temperature for 15 min. The stained cells were observed and photographed using a fluorescent microscope (ZEISS, Germany). The antibodies are listed in Table S3.

### Reactive oxygen species (ROS) analysis

The differently treated cells were digested, washed twice with PBS, and incubated with 10 μM 2ʹ,7ʹ-dichlorofluorescein diacetate (H2DCFDA, Solarbio) in serum-free and antibiotic-free DMEM medium at 37 °C for 30 min in the dark. After washing with PBS, the intracellular ROS levels were then analyzed using a FACSCalibur flow cytometer (BD Biosciences, USA).

### Malondialdehyde (MDA) Assay

The malondialdehyde (MDA) assay kit (Beyotime, China) was applied to detect the level of lipid peroxidation according to the manufacturer’s protocol. In brief, 2× 10^6^ treated or transfected cells were collected and mixed with 150 μl lysis buffer on ice for 30 min. Then 100 μl of sample supernatant was mixed with 200 μl of TBA working solution for 15 min at 100°C, while the residual supernatant of each sample was used for protein concentration detection. After centrifugation, the supernatant was collected, and MDA content was displayed as the absorbance value measured at 532 nm. The MDA concentration was calculated, and the relative content of MDA was finally obtained as the ratio of their MDA concentration to their protein concentration.

### NF-kB reporter assay

The luciferase-based NF-κB reporter vector was obtained from Yeasen Biotech Co., Ltd (China). The NF-κB reporter vector and reference Renilla luciferase vector (Promega, USA) were co-transfected into breast cancer cells at a ratio of 10:1. At 48 h post-transfection, the Dual-Luciferase Reporter Assay System (Promega, USA) was used to measure the luciferase activities with EnSpire™ Multimode Plate Reader (PerkinElmer, USA). The Firefly luciferase activities were normalized with the corresponding Renilla luciferase activities.

### Enzyme-Linked Immunosorbent Assay (ELISA) analysis

The levels of IL-6 secreted by the breast cancer cells in their medium were tested using a CUSABIO^®^ Human IL-6 ELISA kit 6 (CSB-E04638h, Cusabio Biotech Co. Ltd.) based on the manufacturer’s instruction. Briefly, the culture medium of transfected breast cancer cells was replaced by equivalent DMEM without serum 24 h before collection. Then the medium was harvested and the supernatant was obtained by 1,000×g centrifugation for 15 min at 4°C. 100 μl medium supernatant was used for incubation in the ELISA plate, followed by interaction with biotin-labeled IL-6 antibody working solution and HRP-avidin working solution successively. Then, 90 μl transmembrane domain substrate was added into the plate for 15-30min in dark and 50 μl stop solution was subsequently added. Finally, the absorbance was measured at 450 nm by a Microplate Reader (PerkinElmer, USA). The specific concentration value of IL-6 (pg/mL) was calculated based on the standard curve.

### In vivo animal study

Female BALB/c nude mice (4-6 weeks) were purchased from the GemPharmatech Co., Ltd. (Nanjing, China). For subcutaneous inoculation, 1×10^7^ MDA-MB-231 cells stably expressing HSPB1 or control vectors were resuspended in 200 μl PBS and implanted subcutaneously into the right flank regions of the mice. When the subcutaneous tumors reached to an average size of 50 mm^3^, the nude mice from each group were randomly and equally divided into two subgroups, which were treated with 2 mg/kg of DOX or an equal volume of vehicle control via intravenous injection every 3 days for a total of 7 times. Tumor size was measured with calipers every 5 days, and tumor volume was calculated using the formula: volume = length × (width)^2^/2. The mice were sacrificed at the end of experiments, and the subcutaneous tumors were weighed and photographed. For in vivo metastasis assay, 5 × 10^5^ indicated MDA-MB-231 cells were resuspended in 200 μl PBS and intravenously injected into the tail veins of nude mice. 4 weeks later, all the mice were killed and the lungs were harvested and analyzed. Hematoxylin and eosin (H&E) staining was used to confirm the tissue morphology. All animal investigations were approved by the Shandong University Animal Care and Use Committee, and reference number for approval is KYLL-2020(KS)-218.

### Immunohistochemistry (IHC)

The tissues were first fixed in formalin, dehydrated, embedded in paraffin, and then sliced into 4 μm sections. The sections were deparaffinized in xylene and hydrated with gradient alcohol. Citrate buffer was further used for antigen retrieval, and 3% H_2_O_2_ was used to eliminate endogenous peroxidase activity. Sections were then blocked by BSA. Subsequently, the sections were incubated with primary antibody at 4°C overnight. Next day, after washed by PBS, the sections were incubated with peroxidase-conjugated secondary antibody at room temperature for 15 min and stained with diaminobenzidine working solution. Then, the sections were counterstained with hematoxylin, dehydrated with gradient alcohol, mounted, and photographed under the Olympus light microscope. HSPB1 expression was analyzed and scored based on the intensity and the percentage of positively stained tumor cells in the tissue samples, which was evaluated using IHC scores: IHC score = percentage score × intensity score. Percentage staining scores were defined as hereunder mentioned: (i) 0, < 10%; (ii) 1, 10–25%; (iii) 2, 25–50%; (iv) 3, 50–75%; and (v) 4, > 75%; intensity staining scores were divided into four grades as follows: (i) 0, no staining; (ii) 1, light brown; (iii) 2, brown; and (iv) 3, dark brown. Based on above criteria, HSPB1 expression was classified into four grades: (i) negative, IHC score ≤ 3; (ii) weak, IHC score > 3 and ≤ 6; (iii) moderate, IHC > 6 and ≤ 9; and (iii) strong, IHC > 9. The cohort was divided into two groups with HSPB1-high and −low expression by the cut-offs of IHC scores, which was calculated using the receiver operating characteristic (ROC) curves.

### Statistical analysis

Analyses were performed using GraphPad Prism 8.0 Software (La Jolla, CA, USA). The data are presented as mean ± standard deviation (SD) from three independent experiments. P-values were calculated with Student’s t-test or ANOVA test for comparison of two groups or more than two groups. P < 0.05 was considered statistically significant. Survival curves were plotted with the Kaplan–Meier method and compared by the log-rank test. P < 0.05 was considered statistically significant.

## Acknowledgements

This work was supported by National Key Research and Development Program (No. 2020YFA0712400), National Natural Science Foundation of China (No. 82002785; No. 81902695; No. 82002784; No. 81902697), Natural Science Foundation of Shandong Province (No. ZR2020QH258).

## Declaration of competing interest

The authors declare no competing interests.

## Author Contributions

YRL, YJW, and QFY designed the experiments; YRL, YJW, YZ, FZY, DL, YML, YHJ, DWH, ZKW, and XC carried out most of the experiments; BC, WJZ, LJW, TTM, and XLK collected samples; YRL, YJW, FZY, YHJ, and WJZ performed data analysis; YRL, YJW, FZY, and YHJ prepared the figures and tables; YRL, YJW, and QFY wrote and revised the manuscript. All authors read, verified the underlying data and approved the manuscript.

---

**Figure 1-figure supplement 1.**
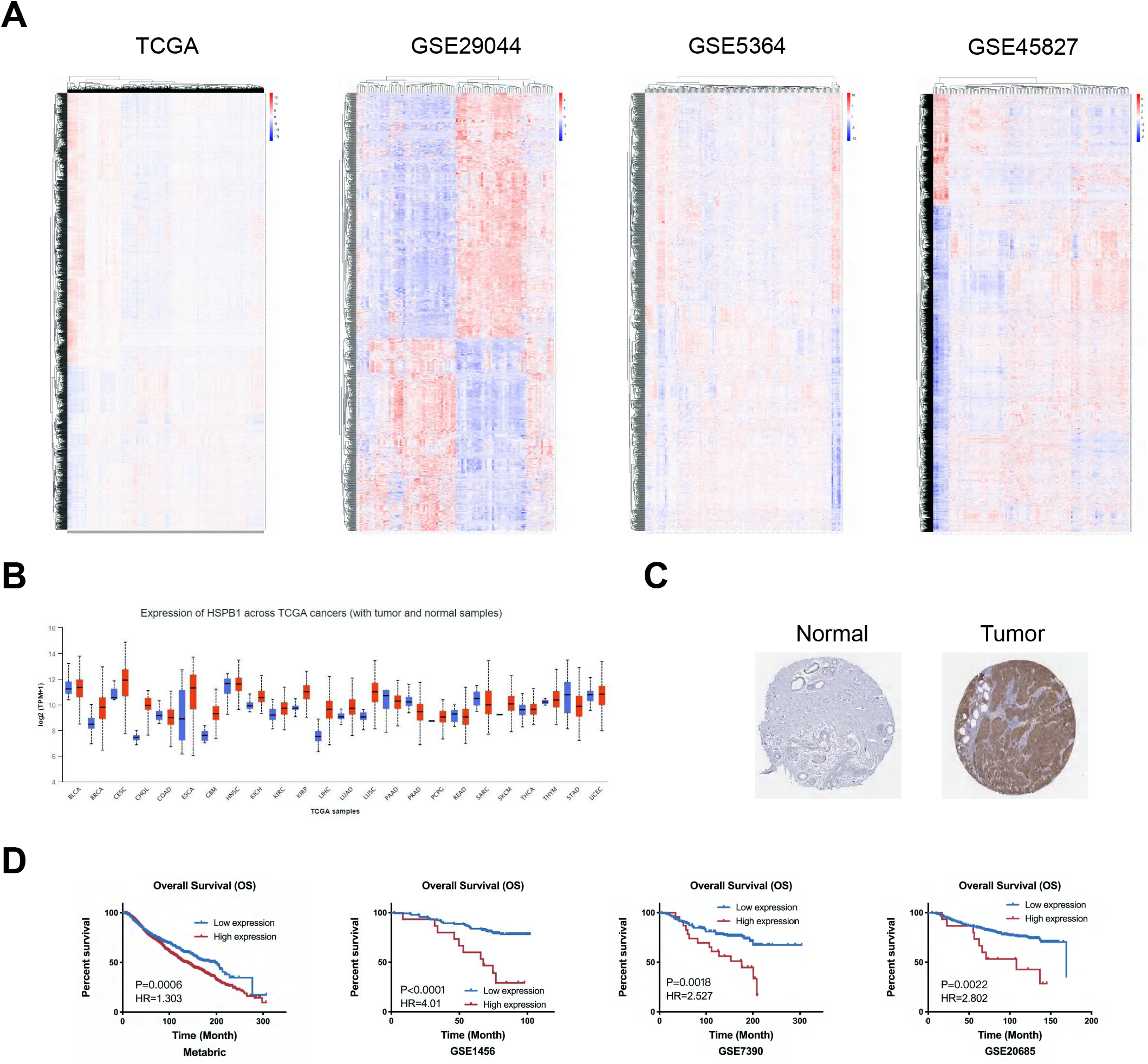
The expression of HSPB1 was upregulated in breast cancer tissues and associated with poor prognosis of breast cancer patients. (A) The heatmaps showing the differentially expressed genes in breast cancer tissues and normal tissues in TCGA and GEO databases. (B) The RNA expression of HSPB1 was upregulated in various cancers according to TCGA database. (C) The protein expression of HSPB1 was increased in breast cancer tissues compared to normal tissues based on the Human Protein Atlas database. (D) High expression of HSPB1 was associated with poor prognosis of breast cancer patients.

**Figure 2-figure supplement 1.**
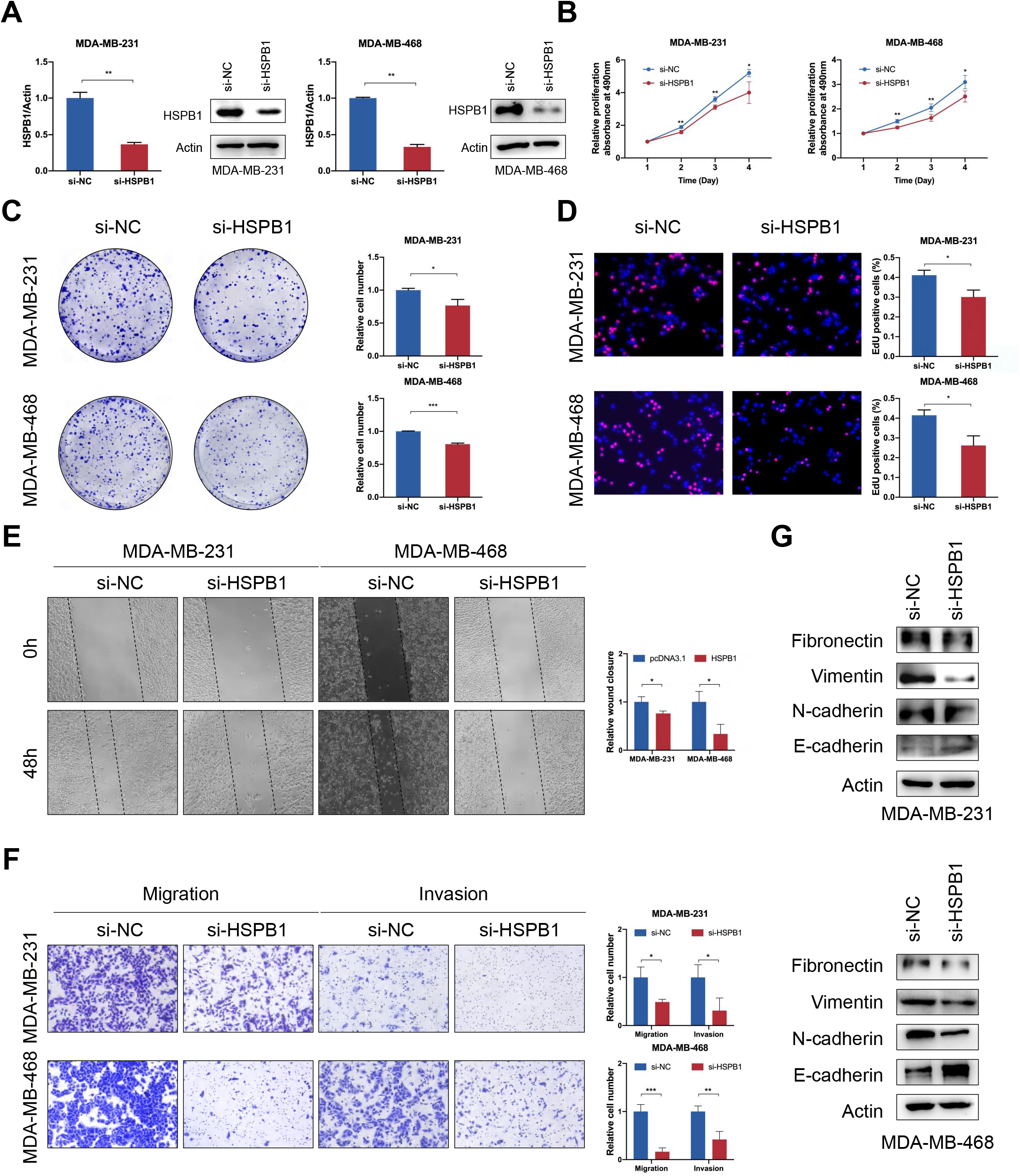
HSPB1 knockdown attenuated breast cancer growth, migration, and invasion in vitro. (A) HSPB1 knockdown efficiency was confirmed by qRT-PCR and western blot in breast cancer cells. (B-D) MTT (B), colony formation (C), and EdU (D) assays were performed to evaluate the effect of HSPB1 knockdown on cell proliferative ability. (E) Wound healing assay was used to evaluate the effect of HSPB1 knockdown on the migration ability of breast cancer cells. (F) The migratory and invasive abilities of HSPB1-knockdown breast cancer cells were assessed by transwell assay. (G) Western blot showed the effect of HSPB1 knockdown on the expression of EMT-related proteins. (* P < 0.05, ** P < 0.01, *** P < 0.001) **Figure 2-figure supplement 1-source data 1** Original western blots for Figure 2-figure supplement 1A. **Figure 2-figure supplement 1-source data 2** Original western blots for Figure 2-figure supplement 1G.

**Figure 3-figure supplement 1.**
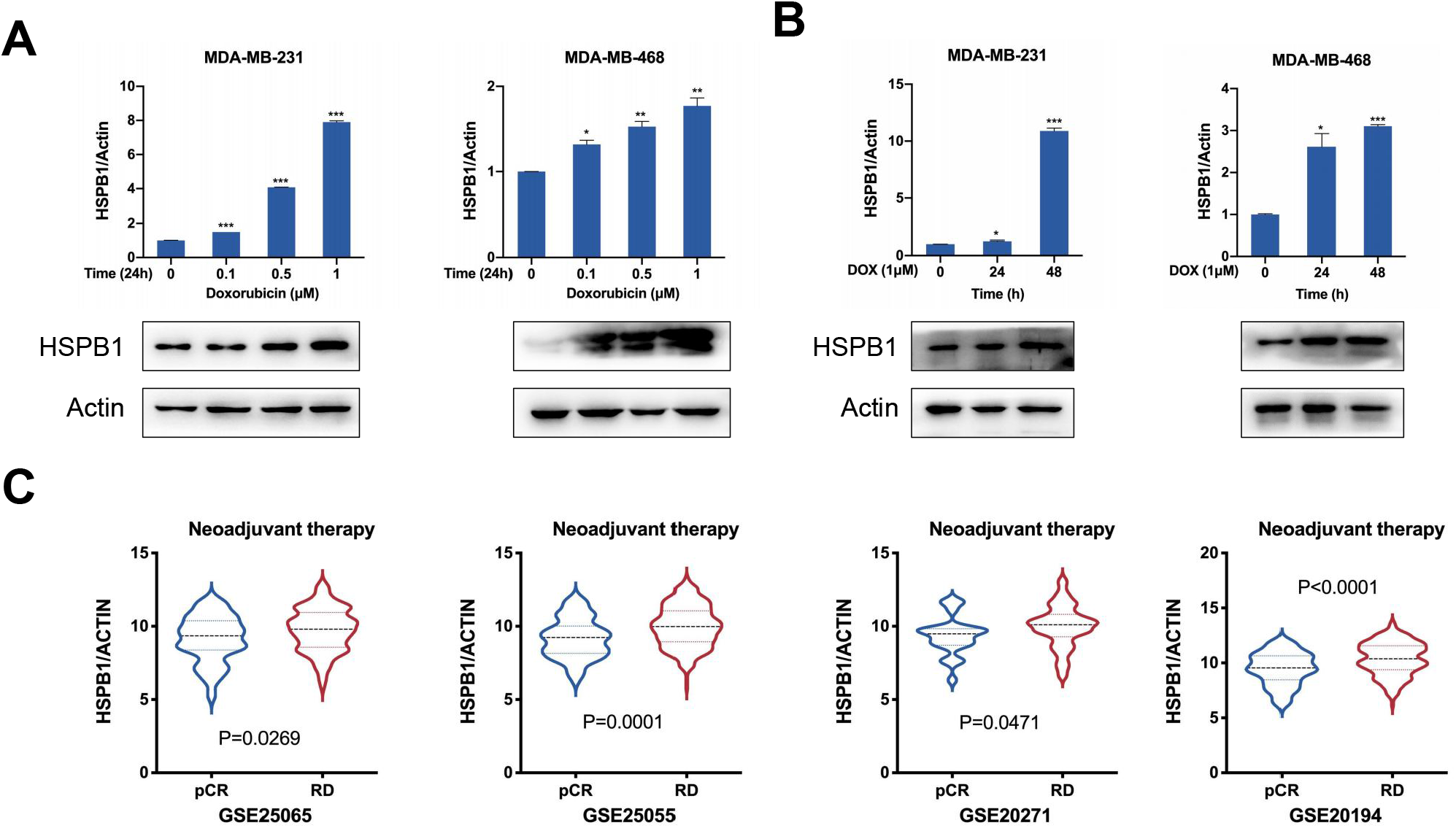
HSPB1 was associated with chemoresistance of breast cancer. (A) MDA-MB-231 and MDA-MB-468 cells were treated with different concentrations of doxorubicin. The RNA and protein expression of HSPB1 was detected. (B) MDA-MB-231 and MDA-MB-468 cells were treated with doxorubicin (1 μM) and collected at the indicated time. The RNA and protein expression of HSPB1 was evaluated. (C) The expression of HSPB1 was upregulated in progressive breast cancer tissues compared to chemo-sensitive tissues. (* P < 0.05, ** P < 0.01, *** P < 0.001) **Figure 3-figure supplement 1-source data 1** Original western blots for Figure 3-figure supplement 1A. **Figure 3-figure supplement 1-source data 2** Original western blots for Figure 3-figure supplement 1B.

**Figure 3-figure supplement 2.**
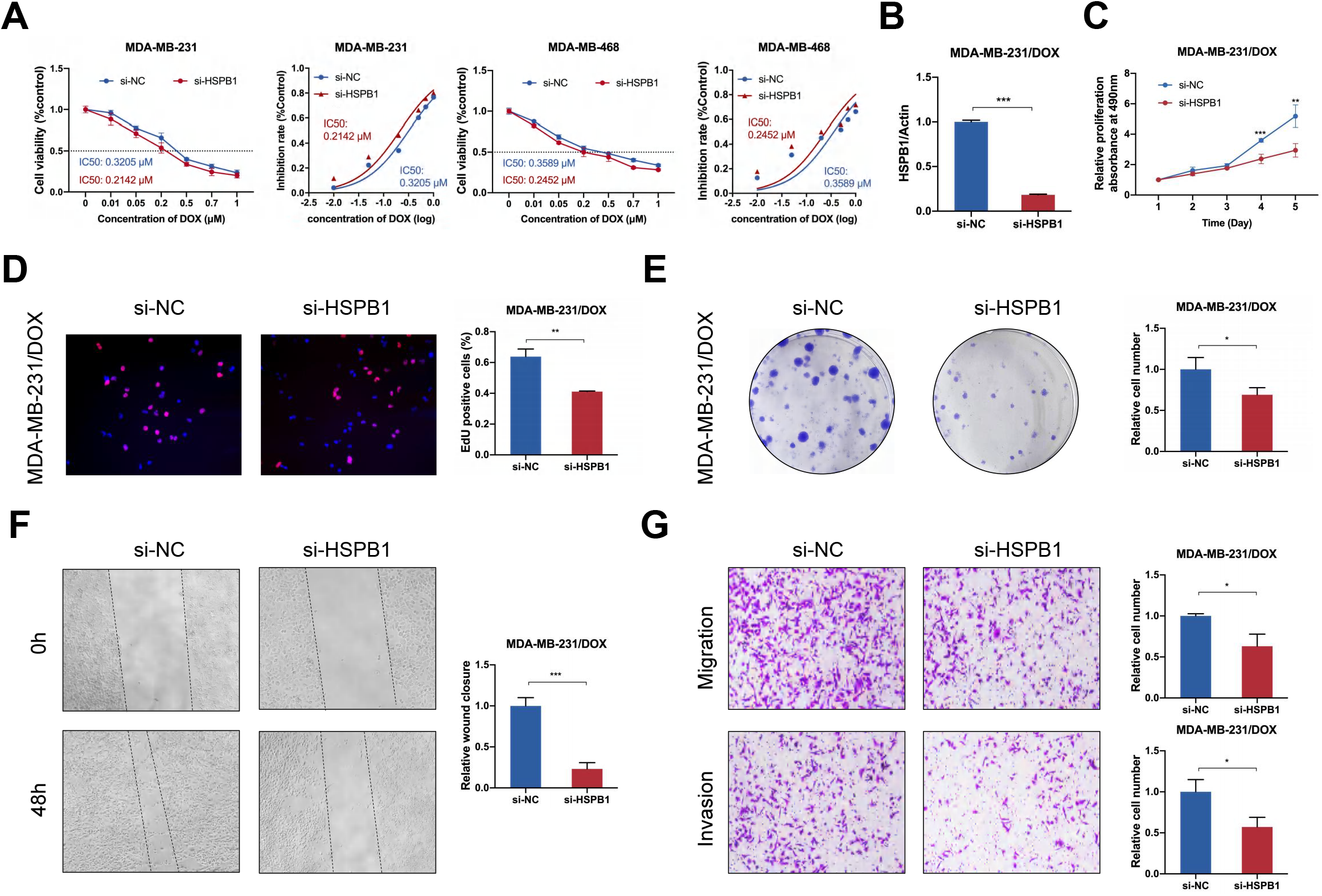
HSPB1 knockdown sensitized breast cancer cells to doxorubicin. (A) Viability of MDA-MB-231 and MDA-MB-468 cells with or without HSPB1 knockdown was analyzed by MTT assay 48 h after treatment with different concentrations of doxorubicin. (B) HSPB1 knockdown efficiency was confirmed by qRT-PCR in MDA-MB-231/DOX cells. (C-E) MTT (C), EdU (D), and colony formation (E) assays were performed to evaluate the effect of HSPB1 knockdown on proliferative ability of MDA-MB-231/DOX cells. (F-G) Wound healing assay (F) and transwell assay (G) were used to assess the migratory and invasive abilities of MDA-MB-231/DOX cells. (* P < 0.05, ** P < 0.01, *** P < 0.001)

**Figure 3-figure supplement 3.**
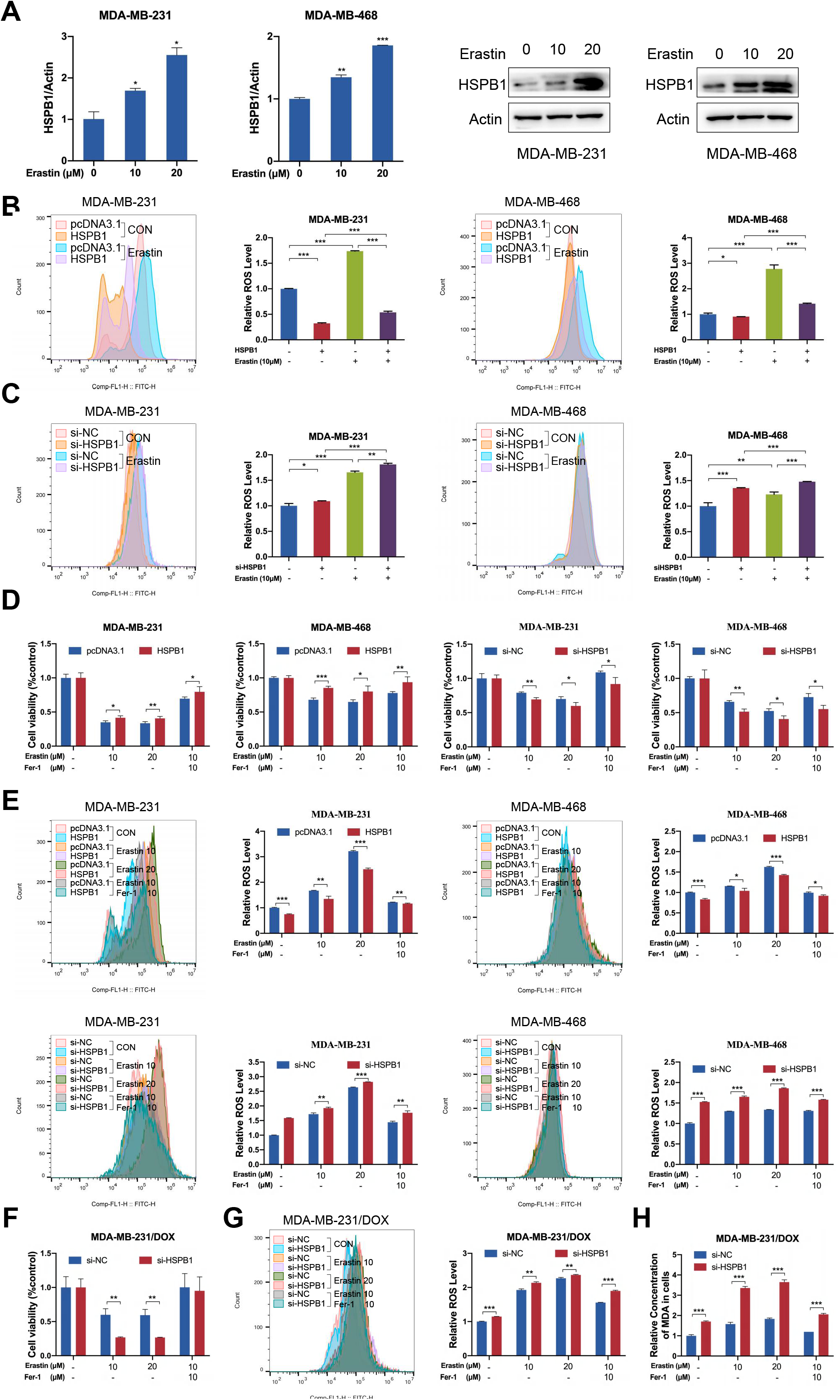
The expression of HSPB1 was associated with ferroptosis. (A) MDA-MB-231 and MDA-MB-468 cells were treated with different concentrations of erastin for 48 h, and RNA and protein expression levels of HSPB1 were assayed by qRT-PCR or western blot. (B) Analysis of lipid ROS in MDA-MB-231 and MDA-MB-468 cells with or without HSPB1 overexpression after treatment with erastin (10 μM, 48 h). (C) Analysis of lipid ROS in MDA-MB-231 and MDA-MB-468 cells with or without HSPB1 knockdown after treatment with erastin (10 μM, 48 h). (D-E) Cell viability (D) and cellular ROS levels (E) in transfected MDA-MB-231 and MDA-MB-468 cells after indicated treatment. (F-H) Cell viability (F), ROS levels (G) and cellular MDA levels (H) were detected in MDA-MB-231/DOX cells with or without HSPB1 knockdown 48 h after treatment with 10 μM or 20 μM erastin plus either DMSO or 10 μM Ferrostatin-1. (* P < 0.05, ** P < 0.01, *** P < 0.001) **Figure 3-figure supplement 3-source data 1** Original western blots for Figure 3-figure supplement 3A.

**Figure 3-figure supplement 4.**
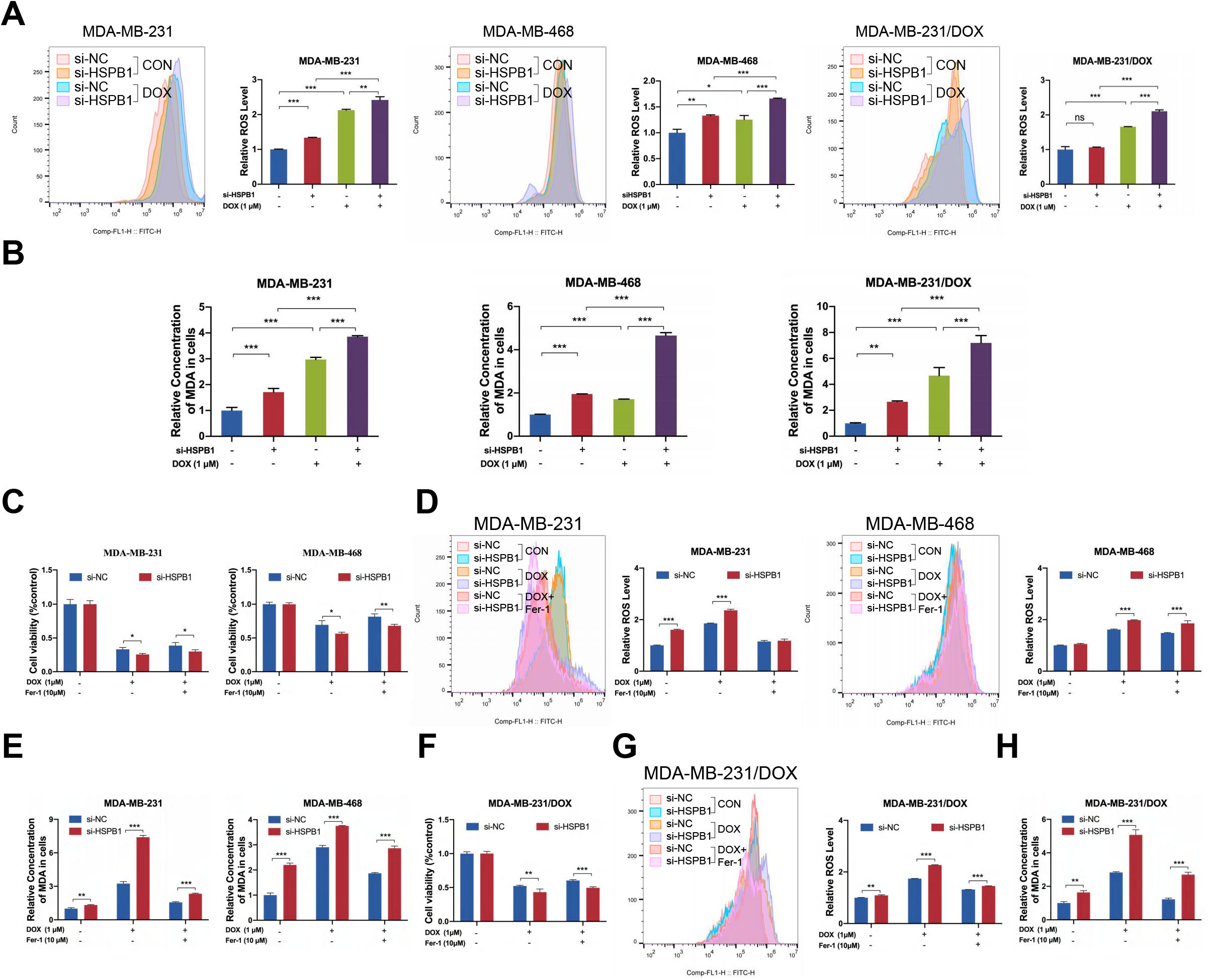
HSPB1 knockdown enhanced doxorubicin-induced ferroptosis in breast cancer cells. (A-B) ROS levels (A) and cellular MDA levels (B) were determined in MDA-MB-231, MDA-MB-468, and MDA-MB-231/DOX cells with or without HSPB1 knockdown 48 h after treatment with 1 μM doxorubicin. (C-E) Cell viability (C), cellular ROS levels (D), and cellular MDA levels (E) in MDA-MB-231 and MDA-MB-468 cells with or without HSPB1 knockdown after indicated treatment. (F-H) Cell viability (F), cellular ROS levels (G), and cellular MDA levels (H) in MDA-MB-231/DOX cells with or without HSPB1 knockdown after treatment with doxorubicin plus either DMSO or Ferrostatin-1. (ns, no significance, * P < 0.05, ** P < 0.01, *** P < 0.001)

**Figure 3-figure supplement 5.**
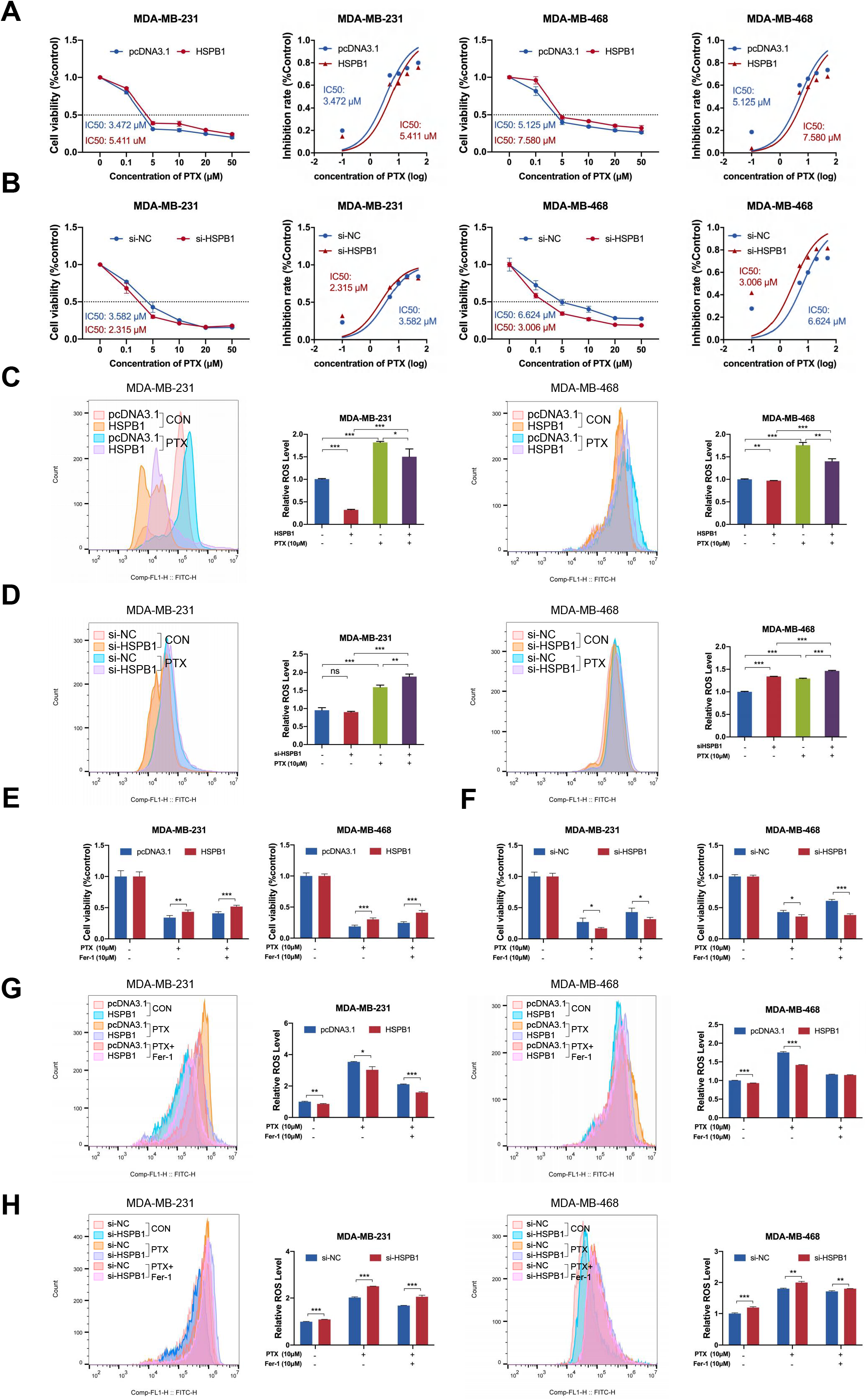
HSPB1 mediated the sensitivity of breast cancer cells to paclitaxel through suppressing ferroptosis. (A) Cell viability of MDA-MB-231 and MDA-MB-468 cells with or without HSPB1 overexpression 48 h after treatment with different concentrations of paclitaxel. (B) Cell viability of MDA-MB-231 and MDA-MB-468 cells with or without HSPB1 knockdown 48 h after treatment with different concentrations of paclitaxel. (C) Cellular ROS levels of MDA-MB-231 and MDA-MB-468 cells with or without HSPB1 overexpression 48 h after treatment with 10 μM paclitaxel. (D) Cellular ROS levels of MDA-MB-231 and MDA-MB-468 cells with or without HSPB1 knockdown 48 h after treatment with 10 μM paclitaxel. (E) Cell viability of MDA-MB-231 and MDA-MB-468 cells with or without HSPB1 overexpression 48 h after indicated treatment. (F) Cell viability of MDA-MB-231 and MDA-MB-468 cells with or without HSPB1 knockdown 48 h after indicated treatment. (G) Cellular ROS levels of MDA-MB-231 and MDA-MB-468 cells with or without HSPB1 overexpression 48 h after indicated treatment. (H) Cellular ROS levels of MDA-MB-231 and MDA-MB-468 cells with or without HSPB1 knockdown 48 h after indicated treatment. (ns, no significance, * P < 0.05, ** P < 0.01, *** P < 0.001)

**Figure 4-figure supplement 1.**
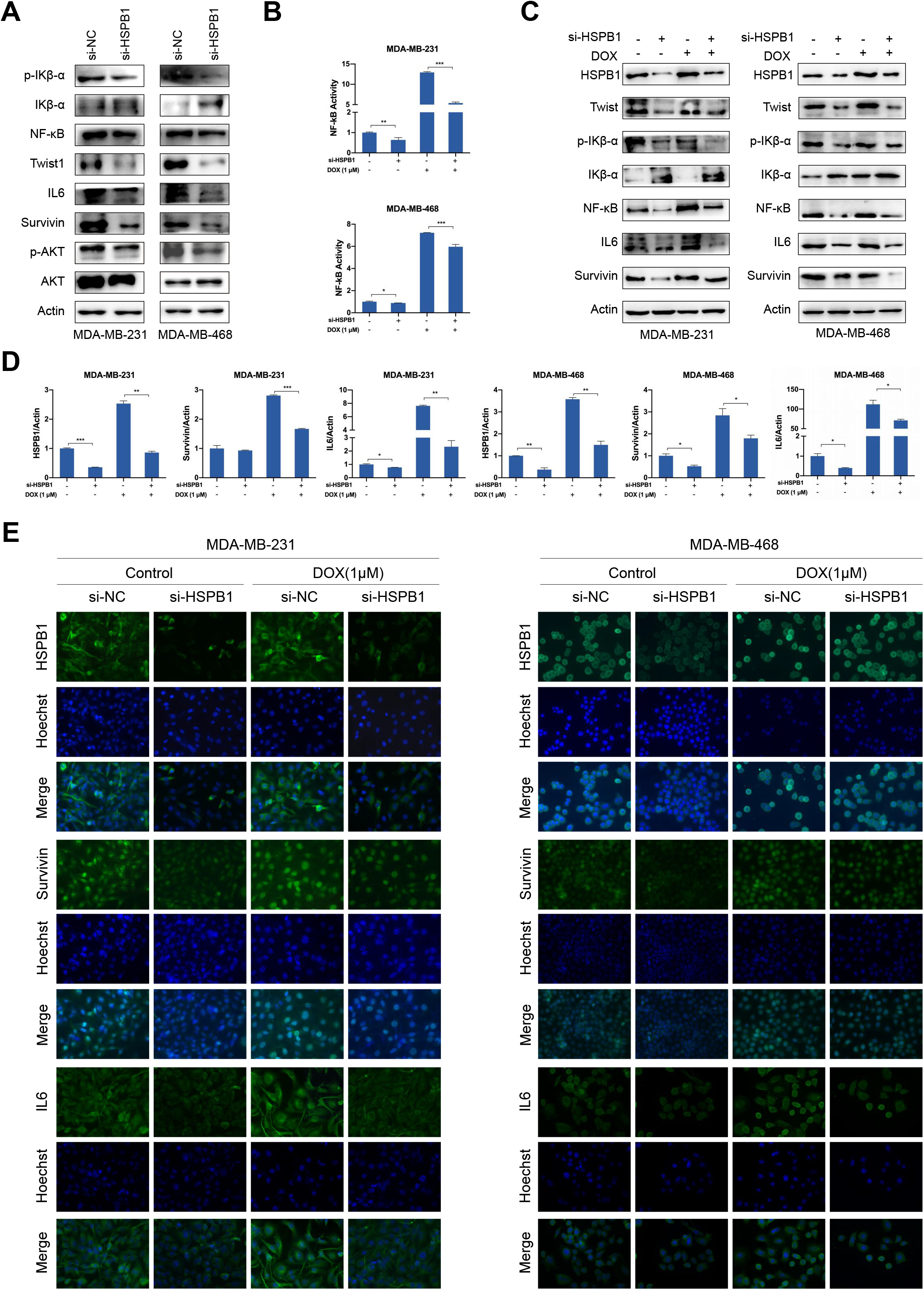
HSPB1 knockdown attenuated activation of NF-κB signaling. (A) Western blot was performed using cell lysates of MDA-MB-231 and MDA-MB-468 cells with or without HSPB1 knockdown. (B-D) NF-κB transcriptional activity was determined by NF-κB activation reporter assay (B), western blot (C), and qRT-PCR (D). (E) After 24h of treatment with 1 μM doxorubicin, immunofluorescence was performed to detect the expression of HSPB1, Survivin, and IL6 in MDA-MB-231 and MDA-MB-468 cells with or without HSPB1 knockdown. (* P < 0.05, ** P < 0.01, *** P < 0.001) **Figure 4-figure supplement 1-source data 1** Original western blots for Figure 4-figure supplement 1A. **Figure 4-figure supplement 1-source data 2** Original western blots for Figure 4-figure supplement 1C.

**Figure 6-figure supplement 1.**
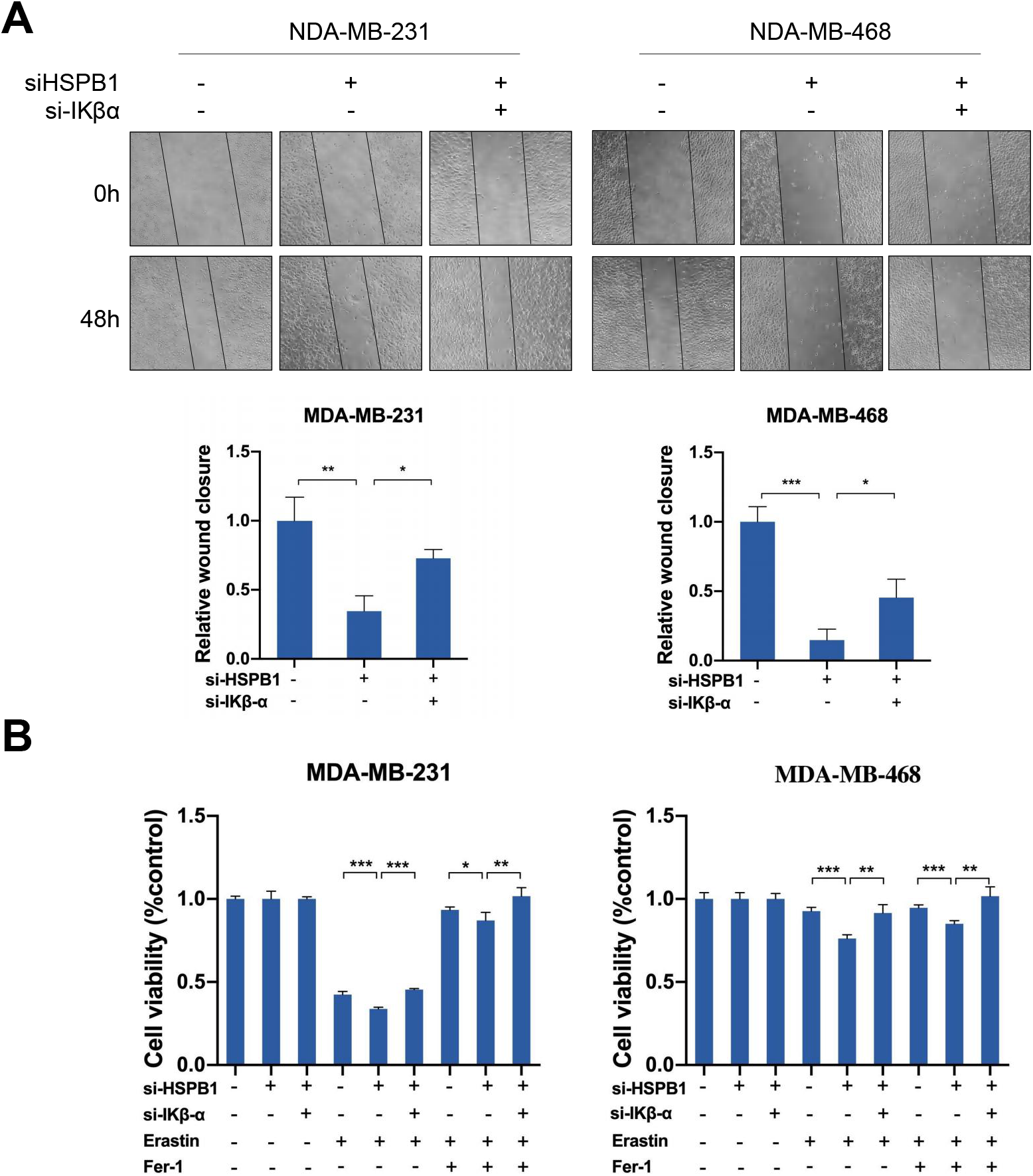
Restoring NF-κB activity partially reversed the inhibited effect of HSPB1 knockdown in breast cancer cells. MDA-MB-231 and MDA-MB-468 cells were transfected with negative control siRNA (Ctrl-siRNA), ctrl-siRNA + si-HSPB1, or si-HSPB1 + si-Ikβ-α for 48 h. (A) Wound healing assay was performed to determined cell migration. (B) The MTT assay was used to analyze the viability of transfected cells following 10 μM erastin plus either DMSO or 10 μM Fer-1 treatment. (* P < 0.05, ** P < 0.01, *** P < 0.001)

**Figure 7-figure supplement 1.**
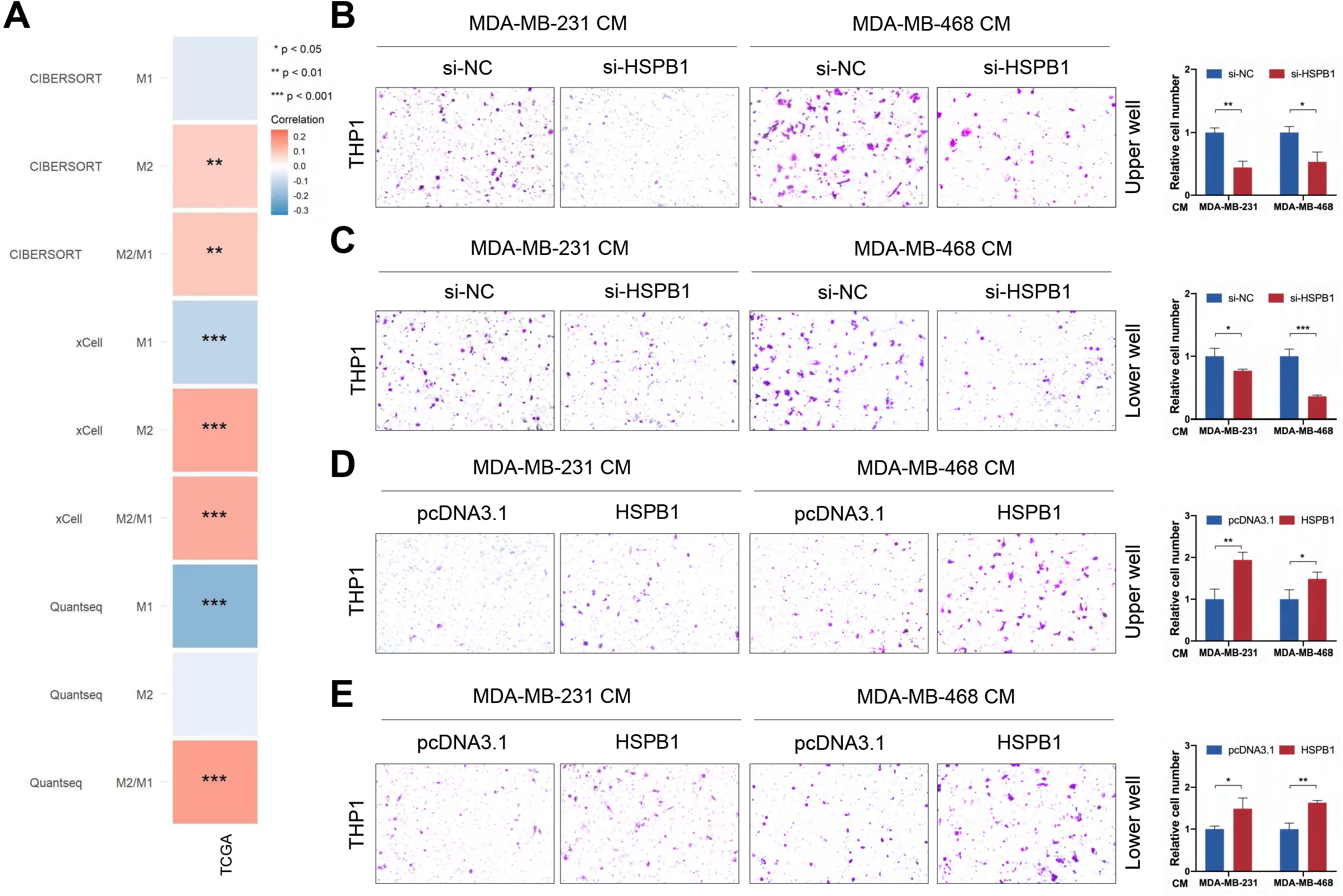
The effect of HSPB1 on the behaviors of THP1. (A) The correlation between HSPB1 expression and the level of macrophage infiltration in tumor microenvironment was analyzed. (B-C) The supernatant of breast cancer cells with or without HSPB1 knockdown was added in the upper chamber or lower chamber, and transwell assay was used to detect the migration of macrophages (B) or chemotaxis (C). (D-E) The conditioned medium from HSPB1 overexpressing cells or control cells was added in the upper chamber or lower chamber, and transwell assay was used to detect the migration of macrophages (D) or chemotaxis (E). (* P < 0.05, ** P < 0.01, *** P < 0.001)

## Supplementary Tables

**Table S1.**
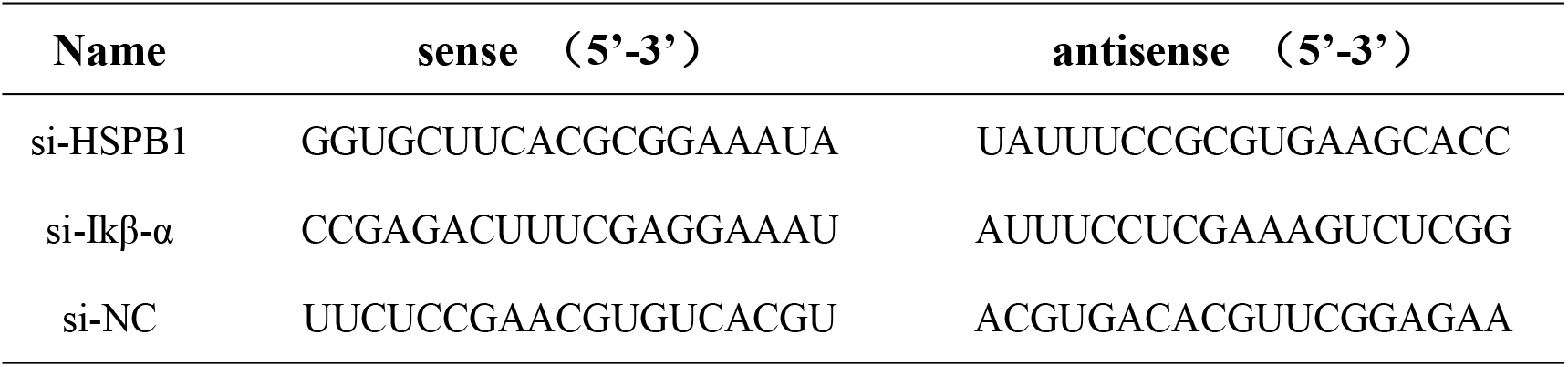
SiRNA used for transfection.

**Table S2.**
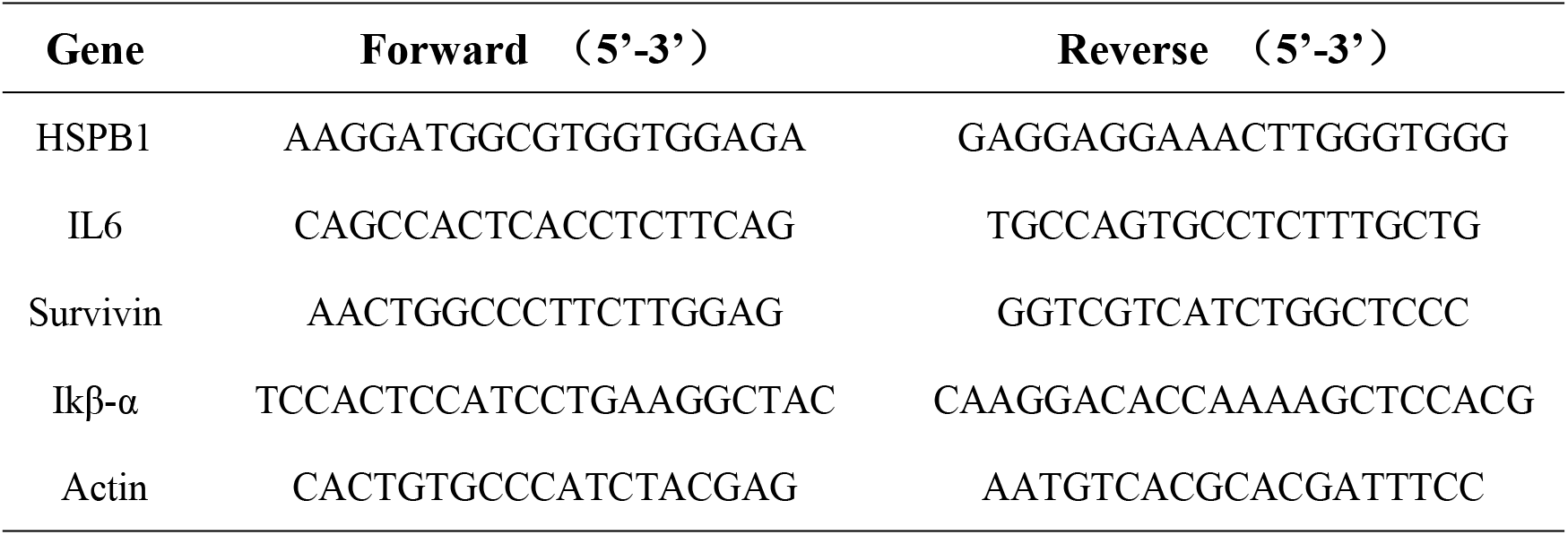
Primers used for qRT-PCR.

**Table S3.** Antibodies used in the experiments.

| Antigen | Supplier | Catalog # | Application |
| --- | --- | --- | --- |
| Actin | Proteintech | 60008-1-Ig | IB (1:2000) |
| Fibronectin | Proteintech | 15613-1-AP | IB (1:1000) |
| N-cadherin | Proteintech | YT2988 | IB (1:1000) |
| E-cadherin | Proteintech | 20874-1-AP | IB (1:1000) |
| Vimentin | Cell Signaling Technology | 5741 | IB (1:1000) |
| HSPB1 | Ptoteintech | 18284-1-AP | IB (1:1000) IF(1:200)<br>IHC(1:200) |
| Twist | Ptoteintech | 25465-1-AP | IB (1:1000) |
| p-IKβ-α | Cell Signaling Technology | 5209S | IB (1:1000) |
| IKβ-α | Ptoteintech | 10268-1-AP | IB (1:1000) IF(1:200)<br>IHC(1:200) |
| NF-κB | Cell Signaling Technology | 8242 | IB (1:1000) IF(1:200)<br>IHC(1:200) |
| IL6 | Ptoteintech | 21865-1-AP | IB (1:1000) IF(1:200)<br>IHC(1:200) |
| Survivin | Ptoteintech | 10508-1-AP | IB (1:1000) IF(1:200)<br>IHC(1:200) |
| p-AKT | Ptoteintech | 28731-1-AP | IB (1:1000) |
| AKT | Ptoteintech | 10176-2-AP | IB (1:1000) |
| Lamin A/C | Ptoteintech | 10298-1-AP | IB (1:1000) |
| Flag | Sigma-Aldrich | F2555 | IB (1:1000) |
| HA | Sigma-Aldrich | H6908 | IB (1:1000) |
| Ki67 | Ptoteintech | 27309-1-AP | IHC(1:500) |
| HRP-anti-mouse | Cell Signaling Technology | 7076 | IB (1:5000) |
| HRP-anti-rabbit | Cell Signaling Technology | 7074 | IB (1:5000) |

## Notes

### Competing Interest Statement

The authors have declared no competing interest.

